# Decrypting the functional design of unmodified translation elongation factor P

**DOI:** 10.1101/2023.09.18.558224

**Authors:** Urte Tomasiunaite, Pavel Kielkowski, Ralph Krafczyk, Ignasi Forné, Axel Imhof, Kirsten Jung

## Abstract

Stalling of ribosomes during polypeptide synthesis due to consecutive proline motifs is a challenge faced by organisms across all kingdoms. To overcome this, bacteria employ translation elongation factor P (EF-P), while archaea and eukaryotes rely on a/eIF5A. Typically, these elongation factors become active only after undergoing post-translational modifications (PTMs) such as ß-lysinylation, (deoxy-)hypusinylation, rhamnosylation, or 5-aminopentanolyation. An exception to this rule is found in EF-P members of the PGKGP-subfamily, which remain unmodified. However, the mechanism behind the ability of certain bacteria to avoid metabolically and energetically costly PTMs, while retaining active EF-P, remains unclear. In this study, we investigated the design principles governing the full functionality of unmodified EF-Ps in *Escherichia coli.* We first screened for naturally unmodified EF-Ps that are active in an *E. coli* reporter strain. We identified EF-P from *Rhodomicrobium vannielii* capable of rescuing the growth deficiencies and changes in the proteome of an *E. coli* Δ*epmA* mutant lacking the gene for the modifying EF-P-(R)-ß-lysine ligase. We then identified specific amino acids in domain I of the unmodified EF-P variant that affected its activity. Ultimately, we transferred these functional properties to other marginally active members of the PGKGP EF-P subfamily, resulting in fully functional unmodified variants in *E. coli*. These results have implications for the improved heterologous expression of polyproline-containing proteins in *E. coli* and offer applications in other bacterial hosts. Understanding the mechanisms that underlie the functionality of unmodified EF-P provides insights into cellular adaptations to optimize protein synthesis.

## Introduction

The translation of mRNAs to proteins at the ribosome is a highly conserved process in all domains of life. The translational efficiency is dynamic and depends on codon bias, tRNA accessibility, and the chemical nature of amino acids to be incorporated into the polypeptide chain ^1^. Among these amino acids, proline presents a unique challenge due to the rigidity of its pyrrolidine ring, making it a poor donor and acceptor during protein translation ^2–5^. Consequently, the presence of consecutive prolines codons leads to a slowdown in protein synthesis and can cause ribosome stalling ^6–9^. Despite the hindrance, polyproline motifs play essential roles in the catalytic activity of enzymes, contribute to protein-protein interactions, and regulate protein copy numbers ^7,10–13^, rendering them indispensable components of the proteome of all organisms.

To facilitate the synthesis of polyproline-containing proteins, organisms from all kingdoms have evolved specialized translation factors. In bacteria, elongation factor P (EF-P) plays a crucial role in this process ^6,7^, while eukaryotes and archaea use initiation factor 5A (e/aIF5A) ^14^. These translation factors bind to the ribosome and enhance peptide bond formation during protein synthesis by stabilizing the P-site tRNA ^15,16^. However, to become fully functional, these translation factors must undergo post-translational modification (PTM) ^6,7,9,14^. Notably, in eukaryotes and archaea e/aIF5A undergoes a unique PTM known as hypusination ^17^. On the other hand, bacteria use diverse and unusual substrates and enzymes to modify their EF-Ps. For instance, in *Escherichia coli* and *Salmonella enterica* a conserved lysine is ß-lysinylated by the EF-P-(R)-ß-lysine ligase (EpmA) and L-lysine 2,3-aminomutase (EpmB), as well as hydroxylated by the EF-P hydroxylase (EpmC) ^18–22^. Other modifications include rhamnosylation of an arginine by the arginine rhamnosyltransferase (EarP) in bacteria like *Pseudomonas putida, Shewanella oneidensis* ^23,24^ and amino-pentanolylation by the EF-P modification enzyme YmfI in *Bacillus subtilis* ^25,26^ **(Fig. 1a)**.

**Figure 1.**
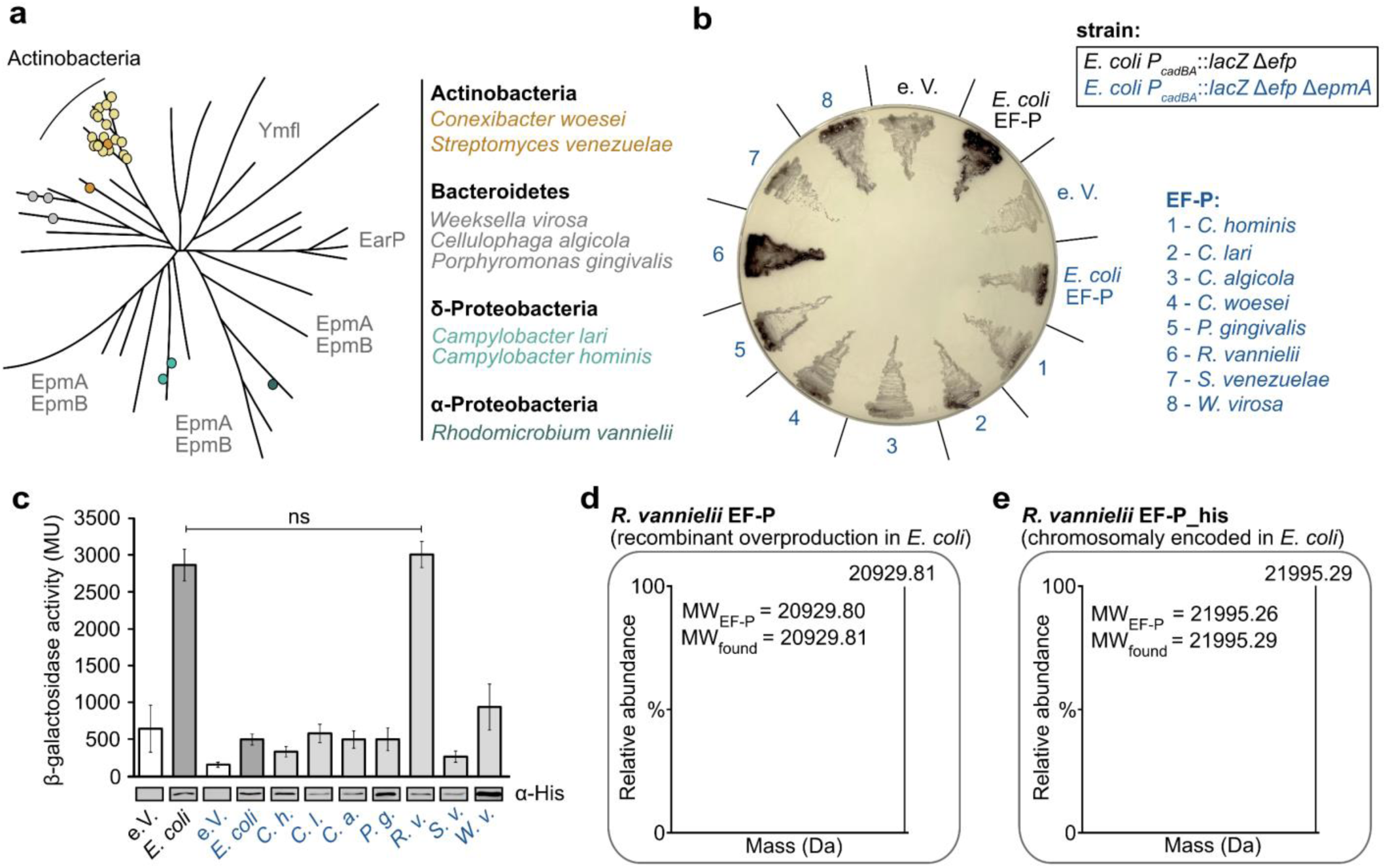
*R. vannielii* EF-P complements Δ*efp E. coli* mutants without the necessity of a PTM. **a** Schematic representation of the phylogenetic tree of bacterial EF-Ps, adapted from Pinheiro *et al.*, 2020 ^27^. PGKGP-subfamily EF-Ps used in this study are marked with coloured dots (Actinobacteria – orange, Bacteroidetes – grey, δ - Proteobacteria – light green, α - Proteobacteria – dark green). Yellow dots represent the remaining PGKGP-type EF-Ps from Actinobacteria. **b**, **c** EF-P activity measurements using the P*_cadBA_*::*lacZ* based reporter assay with S-gal or o-nitrophenyl-β-D-galactopyranoside (*o*-NPG) as substrates for the β-galactosidase activity. Colour code in **c** corresponds to the strains used in **b**. β-galactosidase activities are given in Miller Units (MU). EF-P production was confirmed by Western Blot analysis using antibodies against the His-tag. Error bars indicate 95 % confidence intervals of at least three replicates. Statistics: student’s unpaired two-sided t-test (ns, p = 0.2808). **d** Deconvoluted MS spectra of intact *R. vannielii* EF-P, recombinantly overproduced in *E. coli.* **e** Deconvoluted MS spectra of intact His-tagged *R. vannielii* EF-P (chromosomally encoded), produced *in E. coli*.

Interestingly, there is a group of bacteria, particularly from the Actinobacteria phylum, which possesses functional EF-Ps without the need for any PTM ^27^ (**Fig. 1a**). We have shown that EF-Ps from the genera *Corynebacterium*, *Mycobacterium* and *Streptomyces* function without PTM ^27^. These EF-Ps have the otherwise modified lysine at the tip of a β-hairpin, flanked by two proline residues. It is suggested that the presence of this palindromic sequence Pro-Gly-Lys-Gly-Pro (PGKGP) within the hairpin confers rigidity and enables proper positioning of the protruding lysine 32, thereby stabilizing the acceptor arm of the tRNA. The PGKGP sequence has become a signature motif for this subfamily leading to its designation as the PGKGP-subfamily of EF-P ^27^ **(Fig. 1a).**

While it has been shown that a differentially modified EF-P variant can complement the deletion of *efp* in *E. coli* when co-expressed with the modification machinery ^23^, unmodified EF-P variants were found to be inactive ^27^. This observation suggests that certain features are responsible for conveying functionality without PTM. The current study was conducted to investigate the underlying design principles that enable the evolution of functional EF-Ps without the need for PTMs. We were able to identify three crucial amino acid positions with pivotal roles in maintaining the functionality of unmodified EF-P. Using this knowledge, we then succeeded to engineer a functional unmodified EF-P in the bacterial workhorse *E. coli*. These findings pave the way for novel approaches to optimize synthesis of polyproline-containing proteins at lower metabolic cost in other laboratory and industrially relevant bacteria.

## Results

### 1. Screen for naturally unmodified EF-Ps of the PGKGP-subfamily that are functional in *E. coli*

Previous study has demonstrated that EF-Ps, belonging to the PGKGP-subfamily of various Actinobacteria, are functional without PTM in their respective hosts, but they are non-functional in *E. coli* ^27^. Notably, EF-Ps containing the PGKGP-loop are also found in other bacterial phyla, which do not encode enzymes required for EF-P modification ^27^. This observation has led to the hypothesis that these EF-Ps may also be functional in an unmodified state. Here, we screened several EF-Ps of the PGKGP-subfamily of species from different phyla for their activity in *E. coli*. To this end, we chose *Campylobacter hominis* and *Campylobacter lari* as representatives of the δ-proteobacteria, *Cellulophaga algicola, Porphyromonas gingivalis,* and *Weeksella virosa* as representatives of the Bacteroidetes*, Conexibacter woesei* and *Streptomyces venezuelae* belonging to the Actinobacteria, and *Rhodomicrobium vannielii* as representatives of the α-Proteobacteria (**Fig. 1a**). Despite an overall sequence identity of only 44 % (**Supplementary Table S1**), the PGKGP activation loop remains conserved ^27^.

To elucidate the activity of these EF-Ps in *E. coli*, a well-established reporter system was used ^7^. This reporter system is based on the finding that the transcriptional regulator CadC of *E. coli* is a polyproline protein that requires modified EF-P for its translation. In Δ*efp* mutants, the copy number of CadC is too low to induce the *cadBA* promoter (tested as P*_cadBA_*::*lacZ*). CadC is activated when *E. coli* cells are exposed to low pH in the presence of lysine. To rule out that heterologously produced EF-Ps are modified by the *E. coli*-specific modification system EpmA, a reporter strain with an additional deletion of *epmA* encoding the ß-lysine ligase was used. This reporter strain (*E. coli* P*_cadBA_*::*lacZ* Δ*efp* Δ*epmA)* was transformed with plasmids expressing EF-Ps of different representatives of the PGKGP-subfamily. *E. coli* was grown at pH 5.8 and β-galactosidase activities were determined. On agar plates containing S-Gal and ferric ions, cells producing β-galactosidase can easily be detected by black precipitates. As a control, *E. coli efp* was expressed both in *E. coli* P*_cadBA_*::*lacZ* Δ*efp* and *E. coli* P*_cadBA_*::*lacZ* Δ*efp* Δ*epmA*. Only cells expressing both *efp* and the modification machinery showed black precipitates (**Fig. 1b**). Of the eight selected EF-Ps of the PGKGP-subfamily, only EF-P of *R. vannielii* was active. In contrast to *E. coli* EF-P, *R. vannielii* EF-P produced black precipitates in both reporter strains (**Fig. 1b**). To obtain quantitative results, β-galactosidase activity was determined using a colorimetric assay. *E. coli* EF-P was able to rescue the production of CadC and induce high ß-galactosidase activity in the presence of the PTM machinery (**Fig. 1c**). With the exception of *R. vannielii* EF-P, none of the other EF-Ps was able to complement the *E. coli* Δ*efp* mutant. Remarkably, the activity of the reporter strain producing the *R. vannielii* EF-P was comparable to the activity of the strain with the modified *E. coli* EF-P (**Fig. 1c**).

We then used mass spectrometry-based proteomics (MS) to test whether *R. vannielii* EF-P undergoes post-translational modification in *E. coli*. We first purified recombinantly overproduced *R. vannielii* EF-P using the *E. coli* BL21/pET_SUMO system, which had the advantage that the His-tag could be easily cleaved from the protein after purification. For the detection of the *R. vannielii* EF-P without a His-tag, a polyclonal antibody against this protein was generated (α-625) and used for Western blot analysis. The calculated mass of *R. vannielii* EF-P (20,929.80 Da) was consistent with the measured mass of the intact protein (20,929.81 Da), indicating the absence of a modification (**Fig. 1d, Supplementary Fig. S1a**). This observation was confirmed by analysis of peptides after chymotrypsin digestion (**Supplementary Data_1**). Overproduction of EF-P in *E. coli* may result in a high level of unmodified EF-P ^19^ due to an imbalance between EF-P and its modifying enzymes. To avoid any artefact, we inserted *R. vannielii efp* (encoding a C-terminal *his*-tag) into the *E. coli* genome downstream of the native *efp* promoter and purified this protein (**Supplementary Figures S2a**). The calculated mass of the His-tagged *R. vannielii* EF-P (21,995.26 Da) was consistent with the measured mass of the intact protein (21,995.29 Da) (**Fig. 1e, Supplementary Fig. S1b**), confirming that *R. vannielii* EF-P is unmodified in *E. coli*. This observation was further verified by analyzing the peptides after chymotrypsin digestion (**Supplementary Data_1**). It has to be noted that the sequence available online (GenBank accession number NC_014664.1) for *R. vannielii* EF-P differs slightly from the sequence we determined for the strain used (DSM 162/ATCC 17100) (**Supplementary Figure S3**).

Overall, these data demonstrate that an unmodified EF-P variant can support translation of a polyproline-containing protein in *E. coli*.

### 2. Unmodified *R. vannielii* EF-P is fully functional in *E. coli*

Next, we investigated the functionality of *R. vannielii* EF-P in *E. coli* in more detail. First, we conducted growth tests, including the *E. coli* wild type (wt), the Δ*efp*, Δ*epmA*, Δ*efp* Δ*epmA* mutants, compared to the complemented Δ*efp*::*R. v. efp* and Δ*efp*::*R. v. efp* Δ*epmA* mutants. In the latter two strains, *R. v. efp* is chromosomally encoded under control of the native *E. coli efp* promoter (**Fig. 2a**). After 12 hours of growth in complex (LB) medium, the cell density of the mutants Δ*efp* and Δ*efp* Δ*epmA* was much lower compared to the wt, which is in line with previous observations ^19,28^ (**Fig. 2b**). The mutant producing unmodified *E. coli* EF-P (Δ*epmA)* showed slightly higher cell densities, but still exhibited limited growth compared to the wt (**Fig. 2b**). However, expression of *R. v. efp* in both the Δ*epmA* and the Δ*efp* mutants suspended the growth defect (**Fig. 2b**). To validate growth differences observed with the spot assay, we monitored the growth of the strains in LB medium at 37 °C over time (**Fig. 2c**). The growth defect of the mutant producing the unmodified *E. coli* EF-P (*ΔepmA*) was overcome by the chromosomal insertion of the *R. v efp* (**Fig. 2c**). Only marginal differences in doubling times were detected between the wt and the mutants harboring the *R. v. efp* independent of the presence of EpmA (**Supplementary Table S2**), confirming our hypothesis of a functional *R. v.* EF-P without the need for a PTM in *E. coli*. The observed differences in duplication times between the wt and mutants expressing *R. v. efp* could be attributed to lower copy numbers of *R. v.* EF-P (**Supplementary Figure S2b**).

**Figure 2.**
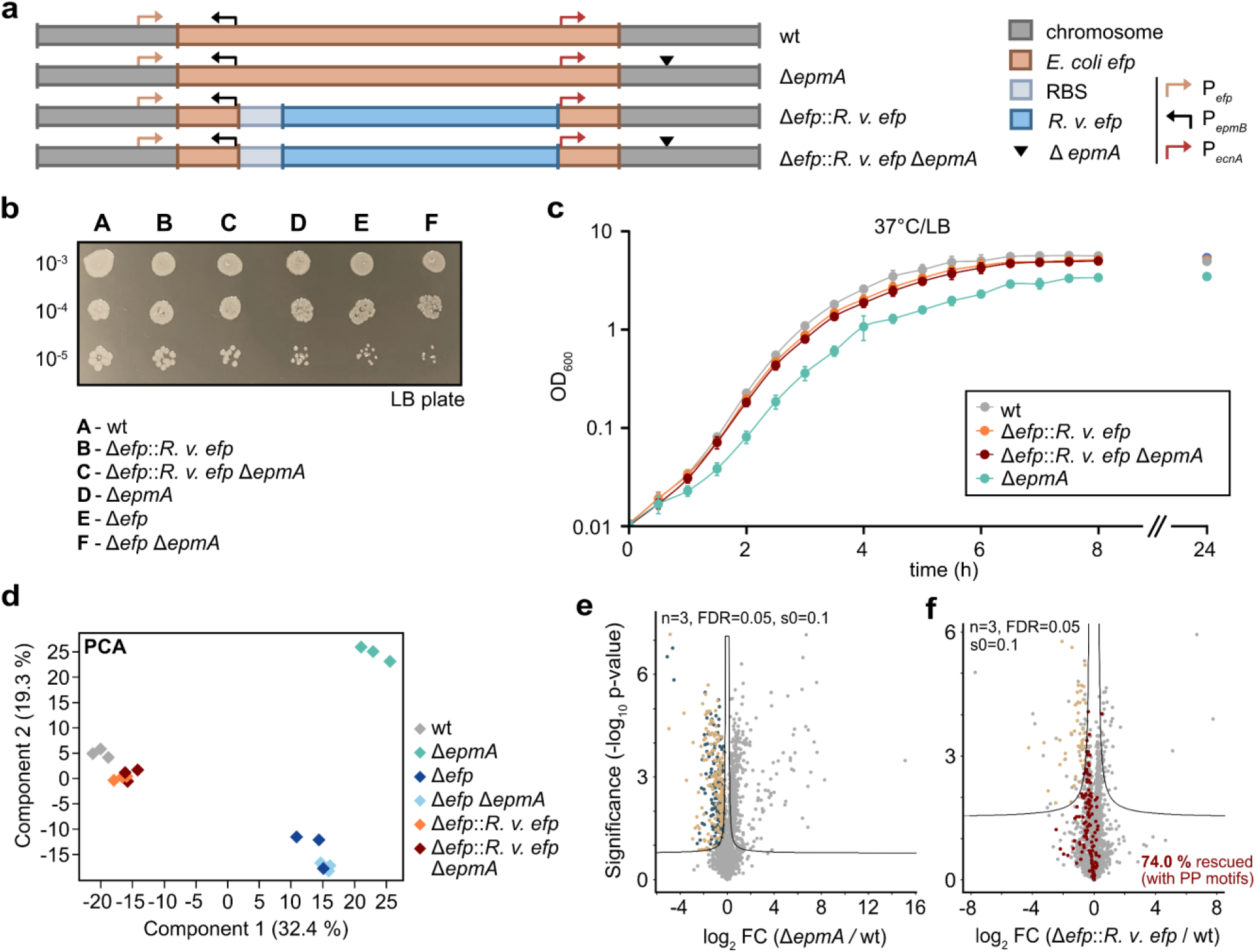
Phenotypic characterization of *E. coli* mutants complemented with *R. vannielii efp.* **a** Schematic overview of *E. coli* BW25113 *efp* deletion mutants with incorporated *R. vannielii efp*. Gene insertions are depicted in coloured boxes, and promoter locations as coloured arrows. **b** Spot assay of the *E. coli* mutants on an agar LB plate after incubation in LB over night at 37°C. **c** Growth of *E. coli* mutants in LB at 37 °C. Data in **c** are mean values with error bars representing the standard deviation (SD) of four independent biological replicates. **d** Proteome comparison of wt, Δ*epmA*, Δ*efp,* Δ*efp* Δ*epmA,* Δ*efp*::*R. v. efp* and Δ*efp*::*R. v. efp* Δ*epmA* mutants using principle component analysis (PCA). Proteomes of three independent biological replicates were analysed and are represented in rhombi with the same colour. **e, f** Volcano plot analysis, highlighting the downregulation of proteins with/without polyproline motifs (PP motifs) in *E. coli* mutant Δ*epmA* (**e**) and their rescue in mutant Δ*efp*::*R. v. efp* (**f**). Downregulated proteins without PP motifs are marked in dark blue dots; with PP motifs - in light brown dots; rescued proteins - in bordeaux red; remaining identified proteins - in grey. The x-axes show the fold change (FC) of the mean value of the log_2_ protein intensity for each protein (LFQ) between two strains. The y-axes show the significance level of the observed difference between the two strains (-log_10_ p-value of the t-test).

The absence of EF-P or its modifying enzymes in *E. coli* is associated with down-regulation of polyproline-containing proteins and alterations of the proteome ^9^. Therefore, we tested whether unmodified *R. vannielii* EF-P inserted into these mutants could restore the wild-type proteome. Cells were grown to late exponential growth phase and prepared for proteomic analysis. The changes in protein levels were determined by LC-MS/MS proteomics using data dependent acquisition (DDA) and label free quantification (LFQ) ^29^. Identified peptide fragments mapped to a total of 2,378 proteins (**Supplementary Data 2**). The differences in proteome comparison between the tested strains can be seen in the principal component analysis (PCA) (**Fig. 2d**). The analyzed proteomes cluster into three populations with large distances between them: (cluster i) wild type together with the two mutants producing *R. vannielii* EF-P (Δ*efp*::*R. v. efp*; Δ*efp*::*R. v. efp* Δ*epmA*), (cluster ii) mutants lacking EF-P (Δ*efp;* Δ*efp* Δ*epmA*), and (cluster iii) the mutant producing unmodified *E. coli* EF-P (Δ*epmA*) (**Fig. 2d**). These data suggest that the proteome of mutants carrying the *R. v.* EF-P show the highest correlation with the wild type *E. coli* strain.

We analyzed the proteomes in more detail with a specific focus on proteins with polyproline motifs. Comparisons between the proteomes of the Δ*epmA* mutant and the wild type (**Fig. 2e),** as well as between the Δ*efp* and Δ*efp* Δ*epmA* mutants and the wild type **(Supplementary Fig. S4a and S4b**) revealed a strong protein scattering in the volcano plot indicating strong differences in the proteomes. Complementation of the Δ*efp* mutant with *R. v. efp* (**Fig. 2f)** or the replacement of *E. c. efp* with *R. v. efp* in the Δ*epmA* mutant (**Supplementary Fig. S4d**) reduced the scattering, suggesting high similarity to the wild-type proteome.

The *E. coli* Δ*efp* and Δ*efp* Δ*epmA* mutants were characterized by down-regulation of proteins with polyproline motifs compared to the wild type (**Supplementary Data 2 and 3**). Replacement of *E. c. efp* with *R. v. efp* allowed translational rescue of 130 (75.6 %) and 152 (78.4 %) polyproline-containing proteins down-regulated in the Δ*efp* and Δ*efp* Δ*epmA* mutants, respectively (**Supplementary Data_2 and 4, Supplementary Fig. S4c and S4d**).

It has been previously reported that unmodified *E. coli* EF-P is still able to rescue the translation of polyproline motifs, but with lower efficiency compared to the modified EF-P ^6,9,30^. Here, we also found that of all the down-regulated polyproline-containing proteins in the Δ*efp* mutant (172 proteins) and the Δ*efp* Δ*epmA* mutant (194 proteins), 47 (27.3 %) and 59 (30.4 %), respectively, were not down-regulated in cells producing unmodified *E. coli* EF-P (Δ*epmA* mutant) (**Supplementary Data_2, 3 and 5)**. Importantly, *R. v.* EF-P was able to outperform the rescue potential of unmodified EF-P in *E. coli* by translationally rescuing 128 (74 %) polyproline-containing proteins (**Fig. 2f; Supplementary Data_2 and 4**).

### 3. Elucidation of the functional design principles of unmodified EF-P in *E. coli*

Our next aim was to uncover the design principles governing the full functionality of unmodified EF-Ps in *E. coli.* Previous reports showed that the bacterial EF-P consists of three domains, forming a shape similar to a tRNA ^31,32^. To identify the essential domains for EF-P, we constructed *E. coli* and *R. vannielii* EF-P hybrids (**Fig. 3a**) and measured their activities using the P*_cadBA_*::*lacZ* reporter system. To determine the boundaries of each domain, we considered predicted secondary structures and 3D protein structures. Linker regions connecting the three domains were selected as fusion points to minimize effects on the EF-P 3D structure (**Supplementary Figure S5a-c**; adapted from Uniprot P0A6N4 and AlphaFold AF-P0A6N4-F1 ^33–35^). The exchange of domains I and II of *E. c.* EF-P with the respective domains of *R. v.* EF-P (H1: *R. v*._*R. v*._*E. c)* more than doubled the activity of *E. coli* EF-P in the Δ*epmA* background (**Fig. 3b**). When only domain I of *E. coli* EF-P was replaced by the corresponding *R. v.* EF-P domain (H2: *R. v*._*E. c*._*E. c)*, the same increase in activity was observed. We concluded that certain amino acids in domain I of *R. vannielii* EF-P are important for its activity in *E. coli*.

**Figure 3.**
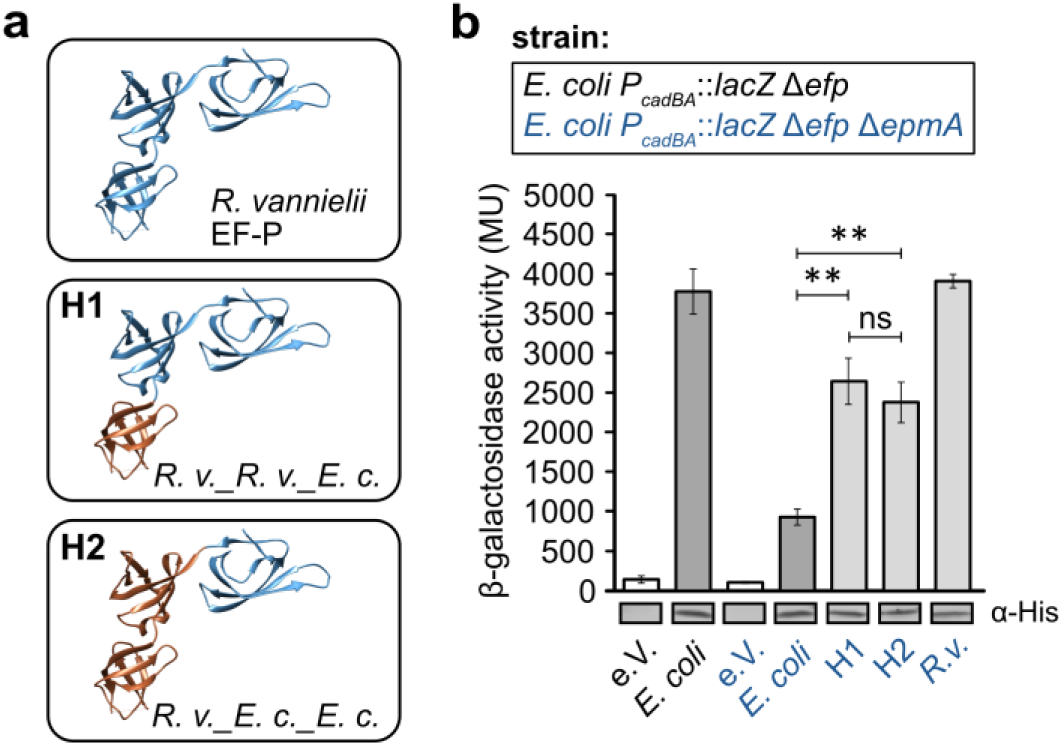
Activity of *E. coli* and *R. vannielii* EF-P hybrids. **a** Schematic overview of the constructed EF-P hybrids H1 (*R. v*._*R. v.*_*E. c*.) and H2 (*R. v*._*E. c.*._*E. c*.). Domains of *R. vannielii* EF-P (*R. v*.) are coloured in blue, those of *E. coli* EF-P (*E. c.*) in brown**. b** Activity measurements of the EF-P hybrids in *E. coli* using the P*_cadBA_*::*lacZ* based reporter assay. β-galactosidase activities are given in Miller Units (MU). EF-P production was confirmed by Western Blot analysis using antibodies against the His-tag. 3D protein structures for simplified EF-P domain representation are taken and modified from PDB 3A5Z. Error bars indicate 95 % confidence intervals of at least three replicates. Statistics: student’s unpaired two-sided t-test (**** p < 0,0001; *** p < 0,001; ** p < 0,01; *p < 0,05; ns p > 0,05). Δ*efp* Δ*epmA* (*E. coli* vs. *R. v.*_H1, ** p = 0.0023; *E. coli* vs. R. v._H2, ** p = 0.0021; *R. v*._H1 vs. *R. v.*_H2, ns p = 0.2023).

To identify these amino acids, we used a synthetic molecular engineering approach, starting with multiple sequence alignments of EF-Ps of the PGKGP-subfamily and *E. coli* (**Fig. 4a**), followed by the construction of EF-P variants using site-directed mutagenesis and screening of those with highest activity using the P*_cadBA_*::*lacZ* reporter assay (**Fig. 4b**). The binding of EF-P to the negatively charged ribosomal parts (rRNA), is mediated mainly by charged amino acid side chains ^15^. Therefore, we focused on amino acids that differ in charge between functional EF-Ps (*E. c.* EF-P and *R. v.* EF-P) and those that are unfunctional (remaining PGKGP-subfamily EF-Ps) in *E. coli*, in particular positions 27, 28, 50, 56, 60, 65 (**Fig. 4a**).

**Figure 4.**
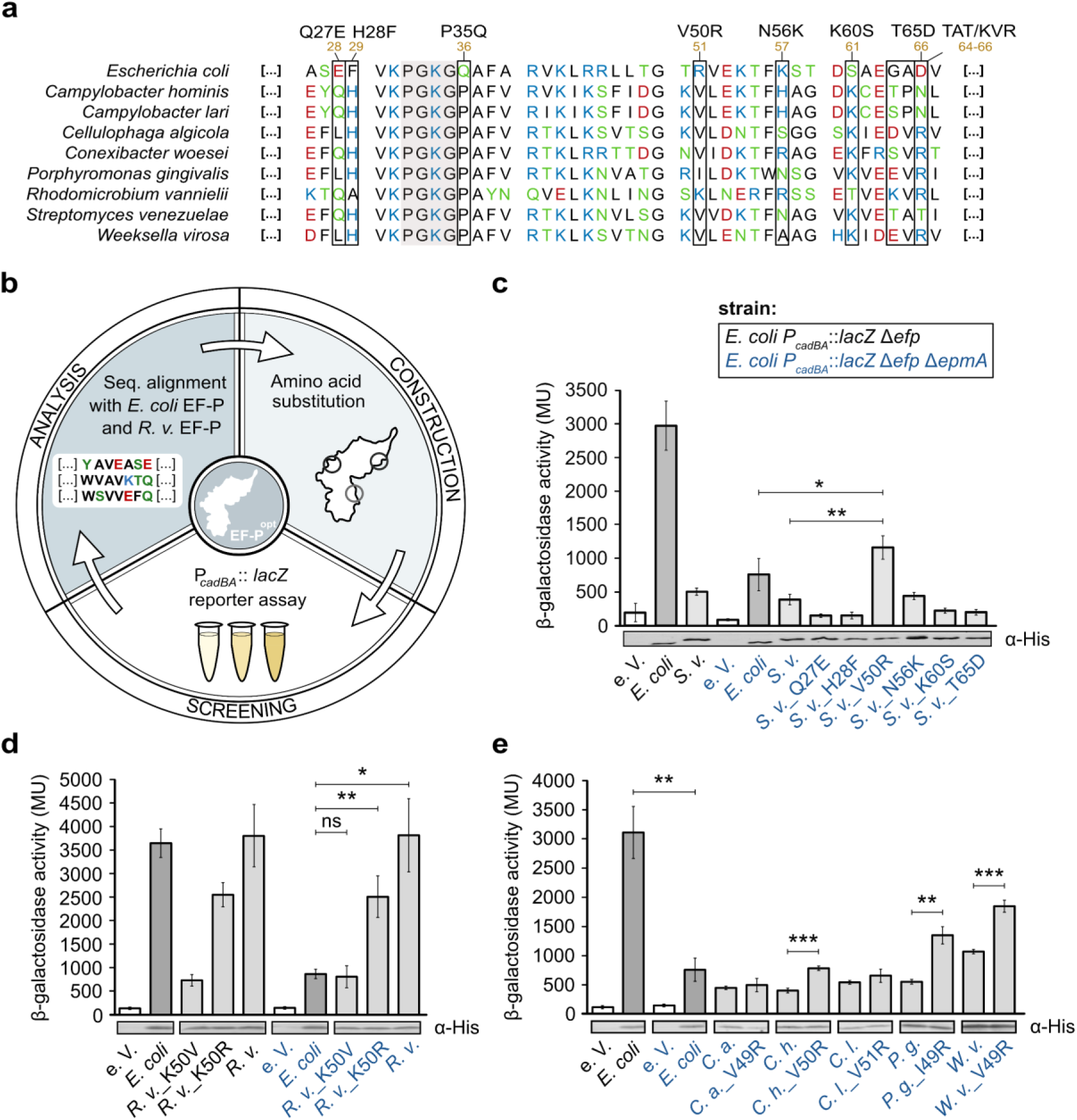
A positively charged amino acid at position 50 has an impact on the activity of PGKGP-subfamily EF-Ps in *E. coli.* **a** Multiple sequence alignment of representative EF-Ps of the PGKGP-subfamily and *E. coli*. Positions selected for substitution are highlighted and numbered according to *S. venezuelae* EF-P. The brown numbers indicate the positions in *E. coli*. **b** Workflow for synthetic engineering of unmodified EF-P. **c**-**e** Activity measurements of *S. venezuelae* EF-P (**c**), *R. vannielii* EF-P (**d**) and PGKGP-subfamily EF-P variants (**e**) in *E. coli* using the P*_cadBA_*::*lacZ* based reporter assay. Colour code in **d** and **e** corresponds to the strains used in **c**. β-galactosidase activities are given in Miller Units (MU). EF-P production was confirmed by Western Blot analysis with antibodies against the His-tag. *C. a. - Cellulophaga algicola, C. h. - Campylobacter hominis, C. l. - Campylobacter lari, E. c. – E. coli, P. g. – Porphyromonas gingivalis, S. v. - Streptomyces venezuelae*, *W. v. - Weeksella virosa*, EF-P^opt^ – optimized EF-P Error bars indicate 95 % confidence intervals of at least three replicates. Statistics: student’s unpaired two-sided t-test (****p < 0,0001; ***p < 0,001; **p < 0,01; *p < 0,05; ns p > 0,05). Δ*efp* Δ*epmA* (*E. coli* vs. *S. v._*V50R, *p = 0.0396; *S. v.* vs. *S. v._*V50R, **p = 0.0035) (**c**); Δ*efp* Δ*epmA* (*E. coli* vs. *R. v._*K50V, ns p = 0.6099; *E. coli* vs. *R. v._*K50R,** p = 0.0092; *E. coli* vs. *R. v.*, * p = 0.0104) (**d**); Δ*efp* Δ*epmA* (*C. h.* vs. *C. h._*V50R, ***p = 0.0001; *P. g.* vs. *P. g._*I49R, **p = 0.0037; *W. v.* vs. *W. v._*V49R, ***p = 0.00099), *E. coli* (in Δ*efp*) vs. *E. coli* (in Δ*efp* Δ*epmA*), ** p = 0.0022 (**e**).

*S. venezuelae* EF-P, a representative of the unmodified actinobacterial EF-Ps with low activity in *E. coli* ^27^, was chosen for molecular engineering (**Fig. 4c**). Out of all substitutions: Q27E, H28F, V50R, N56K, K60S, T65D (numbering according to the *S. venezuelae* EF-P sequence), only the replacement of valine at position 50 to arginine (V50R) significantly increased the activity of *S. venezuelae* EF-P in *E. coli* (**Fig. 4c**). This result suggested that a single amino acid substitution can already impact the overall functionality of EF-P. In the multiple sequence alignment of EF-Ps from the PGKGP-subfamily, we detected a pattern for position 50 (**Fig. 4a**). Strikingly, in almost all EF-P members of the PGKGP-subfamily examined, only uncharged and nonpolar amino acids are present at position 50, with the exception of *R. vannielii* EF-P, which contains a positively charged lysine (K). The EF-P of *E. coli* also has a positively charged amino acid (R, arginine) at this position (**Fig. 4a**). We tested the importance of the positively charged amino acid at position 50. Replacement of lysine by a neutral amino acid (K50V) reduced the activity of *R. vannielii* EF-P in *E. coli* to only 15 %, whereas substitution by another positively charged amino acid, arginine (K50R) allowed 70 % of the original activity (**Fig. 4d**). Based on these results, we investigated the significance of a positively charged amino acid at position 50 in other potentially unmodified EF-Ps from the PGKGP-subfamily. Thus, we constructed EF-P variants of *Cellulophaga algicola*, *Campylobacter hominis*, *Campylobacter lari*, *Conexibacter woesei*, *Porphyromonas gingivalis,* and *Weeksella virosa* EF-Ps with substitutions of valine (V) or isoleucine (I) by positively charged arginine (R) and analyzed their activities using the P*_cadBA_*::*lacZ* reporter assay. Indeed, these amino acid substitutions significantly increased the activity of the EF-P variants of *C. hominis*, *P. gingivalis*, and *W. virosa* (**Fig. 4e**), being in line with results for *S. venezuelae* EF-P (**Fig. 4c**), and highlighting the importance of a positively charged amino acid at position 50 for the activity of unmodified EF-P in *E. coli*. Nevertheless, the introduction of a positively charged arginine (R) at position 50 was not sufficient to increase the activity of EF-P of *C. algicola* and *C. lari* although the proteins were synthesized (**Fig. 4e**).

Overall, these data show that a single amino acid substitution has an effect on the activity of originally non-functional EF-Ps in *E. coli*.

### 4. A few amino acid changes are sufficient to generate a fully functional, unmodified EF-P in *E. coli*

The placement of a positively charged arginine at position 50 resulted in increased activity of *S. venezuelae* EF-P (variant A1) in *E. coli*, although not to the level observed for the modified *E. coli* EF-P (**Figs. 4c, 5a**). Thus, we continued with the synthetic molecular engineering approach (**Fig. 4b**) and investigated other amino acid positions. In particular, we focused on the role of proline 34 (P34), which - together with proline 30 (P30) - is part of the β-hairpin of the PGKGP-subfamily EF-Ps ^27^. Remarkably, substitution of P34 by glutamine (Q) enhanced the activity of the *S. venezuelae* EF-P variant (variant A2) (**Fig. 5a**).

**Figure 5.**
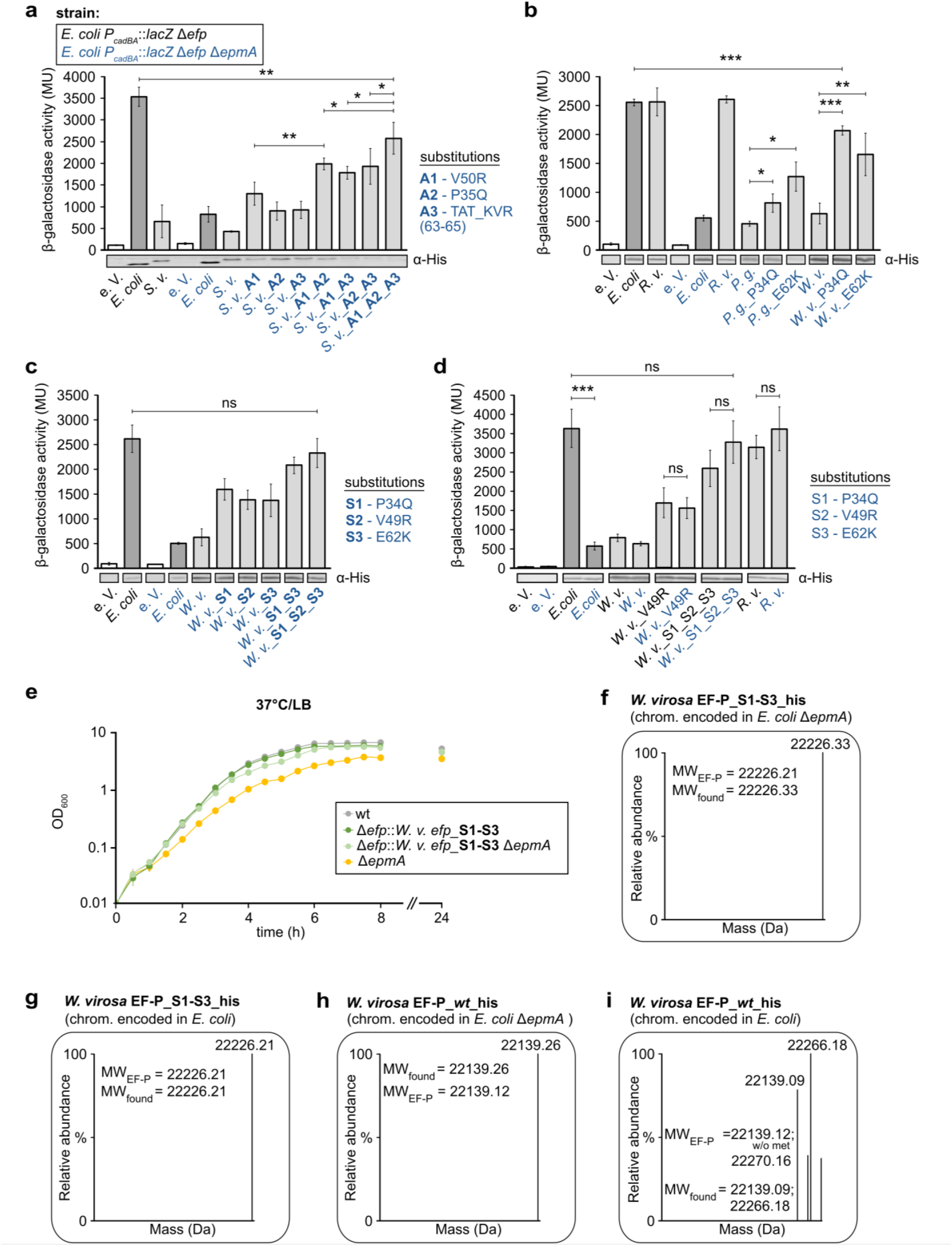
Optimized unmodified EF-P variants are active in *E. coli*. **a**-**c** Activity measurements of EF-P variants from *S. venezuelae* (**a**), *P. gingivalis* (**b**), and *W. virosa* (**b**, **c, d**) in *E. coli* using the P*_cadBA_*::*lacZ* based reporter assay. Colour code in **b**, **c** and **d** corresponds to the strains used in **a**. β-galactosidase activities are given in Miller Units (MU). EF-P production was confirmed by Western Blot analysis using antibodies against the His-tag. **e** Growth of *E. coli* mutants expressing *W. virosa_*S1-S3 *efp* in LB at 37 °C. **f, g** Deconvoluted mass spectra of the chromosomally encoded His-tagged *W. virosa* EF-P _S1-S3, heterologously produced in *E. coli* Δ*epmA* or *E. coli* wt, respectively. It should be noted that the calculated mass of *W. virosa* corresponds to a protein without the first methionine. **h, i** Deconvoluted mass spectra of the chromosomally encoded His-tagged *W. virosa* EF-P wt, heterologously produced in *E. coli* Δ*epmA* or *E. coli* wt, respectively. Calculated masses correspond to *W. virosa* EF-P_wt_6xhis = 22270.16 Da and *W. virosa* EF-P_wt_6xhis without the first methionine = 22139.12 Da. Error bars in the column graphs indicate 95 % confidence intervals of at least three replicates. Data in **e** are mean values with error bars representing the standard deviation (SD) of four independent biological replicates. Statistics: student’s unpaired two-sided t-test (**** p < 0,0001; *** p < 0,001; ** p < 0,01; *p < 0,05; ns p > 0,05). Calculated p-values: Δ*efp* Δ*epmA* (*S. v._*A1 vs. *S. v._*A1_A2, **p = 0.0045; *S. v._*A1_A2 vs. *S. v._*A1_A2_A3, * p = 0.0454; *S. v._*A1_A3 vs. *S. v._*A1_A2_A3, * p = 0.0250; *S. v._*A2_A3 vs. *S. v._*A1_A2_A3, * p = 0.0349), *E. coli* (in Δ*efp*) vs. *S. v._*A1_A2_A3 (in Δ*efp* Δ*epmA*), ** p = 0.0096 (**a**); Δ*efp* Δ*epmA* (*P. g.* vs. *P. g._*P34Q, * p = 0.0244; *P. g.* vs. *P. g._*E62K, * p = 0.0132; *W. v.* vs. *W. v._*P34Q, *** p = 0.0006; *W. v.* vs. *W. v._*E62K, ** p = 0.0094), *E. coli* (in Δ*efp*) vs. *W. v_*P34Q (in Δ*efp* Δ*epmA*), *** p = 0.0005 (**b**); *E. coli* (in Δ*efp*) vs. *W. v_*S1_S2_S3 (in Δ*efp* Δ*epmA*), ns p = 0.1484 (**c**). *E. coli* (in Δ*efp*) vs. *E. coli* (in Δ*efp* Δ*epmA*), *** p = 0.0003; *W. v._*V49R (in Δ*efp*) vs. *W. v._*V49R (in Δ*efp* Δ*epmA*), ns p = 0.4876; *W. v._*S1_S2_S3 (in Δ*efp*) vs. *W. v._*S1_S2_S3 (in Δ*efp* Δ*epmA*), ns p = 0.0506; *R. v.* (in Δ*efp*) vs. *R. v.* (in Δ*efp* Δ*epmA*), ns p = 0.1176; *E. coli* (in Δ*efp*) vs. *W. v.* S1_S2_S3 (in Δ*efp* Δe*pmA*), ns p = 0.2580 (**d**).

Analyzing all positions selected for substitution (**Fig. 4a**), we observed a striking difference between amino acids at position 65 (numbering according to *S. venezuelae* EF-P) in *E. coli* and *R. vannielii*: there is a negative charge in *E. coli* EF-P (aspartic acid, D66) and a positive charge in *R. vannielii* EF-P (arginine, R65). Substitution of T65D did not yield active *S. venezuelae* EF-P in *E. coli* (**Fig. 4c**), which prompted us to replace amino acids of the region 63-65_TAT (T63-A64-T65, threonine – alanine - threonine) with those found in *R. vannielii* EF-P, 63-65_KVR (K63-V64-65R, lysine – valine - arginine). This substitution significantly improved the functionality of *S. venezuelae* EF-P in *E. coli* (variant A3) (**Fig. 5a**).

We further investigated whether the combination of these substitutions could have an additive effect to increase the activity of *S. venezuelae* EF-P (**Fig. 4b**). Indeed, the activity of variant A1_A2 (V50R_P35Q) was significantly higher than that of variant A1 (**Fig. 5a**). Similarly, variants A1_A3 (V50R_63-65_KVR) and A2_A3 (P35Q_63-65_KVR) showed increased activities. Finally, the activity of the variant A1_A2_A3 (P35Q_V50R_63-65_KVR) was almost 6-fold higher than that of the original wild type protein (*S. v.*) (**Fig. 5a**). Nonetheless, the activity of *S. venezuelae* EF-P_A1_A2_A3 remained significantly lower than that of the modified *E. coli* EF-P (**Fig. 5a**).

We found that *S. venezuelae* EF-P_A1_A2_A3 was less well produced compared to *E. coli* EF-P, as indicated by the weaker bands in the Western blots (**Fig. 5a**), which might explain the lower activity. This observation prompted us to search for other EF-P PGKGP-family members that are better expressed in *E. coli.* Since the EF-Ps derived from *P. gingivalis* and *W. virosa* were produced very well in *E. coli* (**Fig. 4e**), we decided to continue synthetic molecular engineering with them. Variants with P34Q or E62K substitutions in both EF-Ps had significantly higher activities than the corresponding native proteins (**Fig. 5b**). It should be noted that in both proteins only the first amino acid (E62K) had to be substituted to obtain the crucial KVR-motif (**Fig. 4a**). Remarkably, the *W. virosa* EF-P variant with all three amino acid substitutions (*W. virosa* EF-P_S1_S2_S3; EF-P_P34Q_V49R_E62K) reached the activity level of the modified *E. coli* EF-P (**Fig. 5c**). The activity of *W. virosa* EF-P_S1-S3 in *E. coli* was also not affected by the presence of the modifying enzyme EpmA (**Fig. 5d**).

To this end, we characterized *E. coli* strains expressing *W. virosa* EF-P instead of their native modified or unmodified EF-P. For this purpose, *E. coli* mutants with chromosomally inserted *W. virosa efp_*S1-S3 were constructed (**Supplementary Fig. S2c**), and Western blot analysis revealed sufficiently high production of this variant (**Supplementary Figure S2d**). *W. virosa* EF-P_S1-S3 was able to rescue the growth phenotypes observed for the Δ*epmA E. coli* mutant independent of the presence of *epmA* (**Fig. 5e**). The chromosomally encoded *W. virosa* EF-P_S1-S3 was purified by affinity chromatography. MS analysis of the intact protein confirmed that this EF-P variant was unmodified. The calculated protein mass of the *W. virosa* EF-P_S1-S3 variant was consistent with the measured intact protein mass, regardless of the presence of EpmA (calculated: 22,226.21 Da; measured: Δ*epmA* 22,226.33 Da; *epmA^+^* 22,226.21 Da) (**Fig. 5f** and **5g; Supplementary Figures S1c** and **S1d**). It should be noted that the measured protein masses are given without the first methionine, a phenomenon frequently observed in previous studies ^36,37^. To test whether the amino acid substitutions affected possible modification events, we also analyzed the intact protein mass of purified *W. virosa* wild type EF-P, and the calculated protein mass (22,139.12 Da) was consistent with the measured intact protein mass, whether or not the cells expressed EpmA (Δ*epmA:* 22,139.26 Da; *epmA^+^*: 22,139.09 Da) (**Fig. 5h** and **5i**; **Supplementary Figures S1e** and **S1f**).

In summary, these data show that three amino acid substitutions are sufficient to convert an inactive EF-P into a fully functional unmodified variant. Moreover, an *E. coli* mutant is now available as a host for heterologous production of polyproline-containing proteins with lower metabolic and energy costs, since EF-P requires no longer PTM.

## Discussion

For optimal synthesis of proteins with polyproline stretches, the EF-P of *E. coli* requires a PTM catalyzed by the modifying enzymes EpmA, EpmB and EpmC ^18–21^. EpmB converts the precursor (S)-α-lysine to (R)-β-lysine, which is then ligated by EpmA in an ATP-dependent condensation reaction to the ε-amino group of Lys34 in EF-P. EpmC subsequently catalyzes the hydroxylation of Lys34. Except for hydroxylation, none of the modification steps can be bypassed without noticeable effects on translational rescue of PP-proteins. Even modification of EF-P with α-lysine instead of ß-lysine decreases its activity ^38^. Notably, the absence of modification in EF-P impairs the translation of polyproline-containing proteins ^9^. This study revealed that only one third of polyproline-containing proteins found to be downregulated in the Δ*efp* mutant, were rescued by unmodified E-FP (Δ*epmA* mutant) (**Supplementary Data_2, 3 and 5**). Thus, while the lack of PTM does not completely prevent EF-P from its original mode of action, it impacts its efficiency. The PTM of *E. coli* EF-P catalyzed by EpmA, EpmB and EpmC is associated with metabolic (ß-lysine) and energetic (ATP) costs for *E. coli*.

The relatively high abundance of diproline-motifs (XPPX)-containing proteins in *E. coli* (2,101 motifs, 0.49 motif/protein) ^13^, shown to cause ribosome stalling ^9^, underscores the need for an EF-P with maximal functionality to maintain proteome homeostasis, especially at high growth rates ^28^. Many representatives of Actinobacteria have an even higher number of XPPX motifs within their proteomes^27^. In *Streptomyces coelicolor* and *Mycobacterium tuberculosis,* the number of XPPX motifs even surpasses the number of encoded proteins (1.08 motif/protein and 1.17 motif/protein, respectively) ^27^. Accordingly, one would expect PTMs to be crucial in these bacteria. However, the actinobacterial EF-Ps are not modified ^27^. Both *S. coelicolor* and *M. tuberculosis* grow very slowly ^27,39–41^. In these slowly growing bacteria, an unmodified, and therefore metabolically and energetically less demanding EF-P could provide a selective advantage when coping with the production of a large number of polyproline-containing proteins.

However, actinobacterial unmodified EF-Ps could not rescue the translation of PP proteins in *E. coli* ^27^. To find out why certain EF-Ps exhibit functionality in their native host without PTM, but are non-functional in *E. coli*, we analyzed the activity of EF-Ps belonging to the PGKGP-subfamily from different phyla (**Fig. 1a**). We found that of all eight tested PGKGP-subfamily EF-Ps, the EF-P from *R. vannielii* was able to rescue the translation of polyproline-containing proteins in *E. coli* (**Fig. 1b, c**). Importantly, the EF-P from *R. vannielii* retained its functionality in an unmodified state in *E. coli* (**Fig. 1d, e**). *E. coli* strains expressing *R. vannielii efp* rescued growth defects observed in *E. coli* mutants lacking *efp* or *epmA*. Moreover, the proteome profiles of *R. vannielii* EF-P-producing strains closely resemble those of the wild type (**Fig. 2b-f**).

EF-Ps have conserved regions required for ribosomal contact and adopt a tRNA-like structure, hinting at the evolutionary origin of these proteins ^31,42^. The analogy of EF-P to tRNA, which is one of the building blocks of the translational machinery, likely increased the ability of this protein to bind to the ribosome ^42^ **(Supplementary Figure S6a**). Therefore, we hypothesized that inactive EF-Ps might essentially be functional in *E. coli* but hindered from contributing to the translation process by week binding to the ribosome or insufficient contact to the tRNA. To investigate this, we developed a synthetic molecular engineering workflow involving the substitution of amino acids based on the sequence of the functional *R. vannielii* EF-P (**Fig. 4b**). By replacing single amino acids at positions 35, 50 and 63-65 (numbered according to *S. venezuelae* EF-P, **Fig. 4a**), we successfully converted initially nonfunctional EF-Ps of *S. venezuelae*, *P. gingivalis* and *W. virosa* into functional ones in *E. coli* (**Fig. 4c, e**; **Fig. 5a-d**). Previous studies have reported the importance of certain amino acids at position 35 for EF-P activity. Pinheiro *et al.* showed that a proline at position 34 (position 35 in *S. venezuelae*) is necessary for EF-P to function with highest activity in *Corynebacterium glutamicum*, as substitutions to alanine, glutamine, glycine and asparagine decreased activity of the corresponding EF-P variants ^27^. In *E. coli*, a glutamine (Q) at the corresponding position, has been shown to be required for proper function of EF-P ^30^. These observations are in line with our results, as substitution of this proline to glutamine significantly increased the activity of the corresponding *S. venezuelae*, *P. gingivalis* and *W. virosa* variants in *E. coli* (**Fig. 4a-c**). It is suggested that the amino acid at position 35 is essential for the correct positioning of the conserved lysine (either modified or non-modified) in the ß-hairpin to get contact with P-site tRNA ^15,27^. The presence of a glutamine at this particular position could be important to allow contacts with the *E. coli* ribosome (23*S* rRNA) and consequently stabilization and/or orientation of the ß-hairpin with the essential lysine at the tip in the absence of a modification **(Supplementary Figure S6b**).

Another interesting finding was that substitution of amino acids at position 50 into an arginine (R) resulted in an increase in activity for all tested variants (**Fig. 4e**). This positively charged amino acid is predicted to be in close contact with the ribosome ^15,16,42^ and therefore important for the corresponding EF-P to establish contacts with the ribosome (**Supplementary Figure S6c**). We noted that a positively charged amino acid at position 50 is also found in some EF-P orthologs, the eukaryotic/archaeal initiation factors 5A (e/aIF5A) ^31^(: Sequence alignment_Fig.4). Finally, the substitution of GAD (positions 64_66 in *E. coli*) against KVR had a major impact on the functionality of the unmodified EF-P variants and might be explained by a stabilization of the interaction between EF-P and the P-site t-RNA ^15^ (**Supplementary Figure S6d**).

In conclusion, bacteria, archaea and eukaryotes have evolved different ways to cope with translation of polyproline-containing proteins. There are functional EF-Ps that do not necessitate modification, as well as those requiring PTMs. Our study reveals that EF-Ps with minimal activity can be rendered functional by altering distinct amino acids, presumably influencing the interaction between EF-P, the ribosome and the t-RNA in the P-site. During evolution, similar activation has been achieved by recruiting post-translational modifications. This aligns with a previous study that demonstrated the transformation of the inactive EF-P from *Shewanella oneidensis* in *E. coli* into an active and rhamnosylated form after co-production of the glycosyltransferase EarP ^23^.

The modification of EF-P with ß-lysine in *E. coli* requires not only a substrate but also energy-intense enzymes to catalyze this process. Here we describe the first EF-P that is functional in *E. coli* without the need for a PTM, making it attractive for future studies aiming to reallocate cellular energy for other energy-consuming processes in bacteria. An optimally functioning EF-P is important for proteome homeostasis and bacterial virulence ^43,44^. Thus, decrypting the functional design of unmodified EF-P in *E. coli* not only directs protein production in a resource-efficient manner for scientific and industrial applications but also extends the fundamental understanding of the functional principles of this translation factor.

## Material and Methods

### Lead contact and material availability

Further information and requests for resources and reagents should be directed to and will be fulfilled by the Lead Contact, Kirsten Jung. All unique reagents generated in this study are available from the Lead Contact with a completed materials transfer agreement.

### Bacterial growth conditions

*E. coli* was cultivated in lysogeny broth (LB) supplemented with antibiotics under agitation (750 rpm) at 37 °C. Bacterial growth experiments were conducted in 50 mL glass flasks filled with 15 mL LB. For the spot assay, overnight cultures were resuspended in LB to an optical density (600 nm) (OD_600_) of 0.01, spotted in dilutions (10^-3^ to 10^-5^) on LB plates and grown for 18 h at 37°C. For CadC production, cells were grown in buffered LB at pH 5.8 ^7^. For EF-P overproduction, growth media were supplemented with L-arabinose [0.2 % (w/v)] and chloramphenicol. Antibiotic concentrations used in this study: 34 μg/mL chloramphenicol, 50 µg/mL kanamycin sulphate.

### Plasmid and bacterial strain construction

All primers, plasmids and strains used and constructed in this study are shown in **Supplementary Data_6**. All PCR reactions were conducted using the Q5 polymerase (New England BioLabs) according to manufacturer’s instructions. Standard DNA restrictions were performed in rCutSmart buffer and the fragments were ligated into the corresponding vectors using the T4 ligase (New England BioLabs). e*fp* genes from various species including a sequence coding for a C-terminal 6xHis tag were cloned in pBAD33. Plasmids were isolated using Hi Yield® Plasmid Mini Kit (Sued Laborbedarf), whereas PCR fragments from the agarose gel were purified using the High-Yield PCR Cleanup and Gel Extraction Kit (New England BioLabs). Site-specific mutagenesis of *efp* was carried out with the Q5® Site-Directed Mutagenesis Kit (New England BioLabs) according to manufacturer’s instructions.

*E. coli* mutants expressing *efp* variants were constructed using double homologous recombination with neomycin acetyltransferase and SacB as the selection or counterselection markers, respectively ^45^. Overlapping regions of *E. coli efp* with the promoter of its modification enzyme *epmB* and the lipoprotein entericidin A (*ecnA*) were considered by keeping the 5’ and 3’ gene regions of *E. coli efp* present in the chromosome ^46^ (**Fig. 2a, Supplementary Figure S2a, c).** All nucleotide and protein sequences were analyzed using CLC Main Workbench 8.1.2 (Qiagen).

### β-Galactosidase activity assays

Reporter strains MG1655 Δ*lacZ* P*_cadBA_*::*lacZ* Δ*efp* and MG1655 Δ*lacZ* P*_cadBA_*::*lacZ* Δ*efp* Δ*epmA* transformed with plasmids expressing *efp* and its variants were inoculated in 1.8 mL buffered LB pH 5.8 (91.5 mM KH_2_PO_4_; 8.5 mM K_2_HPO_4_; 0.2 % (w/v) arabinose; chloramphenicol) and microaerobically grown under agitation (Thermomixer Comfort; 700 rpm) at 37 °C overnight. The overnight culture was split into three tubes to measure optical density, ß-galactosidase activity and detect EF-P in a Western Blot. Activity measurements were conducted according to the protocol described previously ^7,27^. The measured β-galactosidase activity is given in Miller Units (MU), calculated according to Miller *et al.*, 1992 ^47^.

### Protein purification

For the recombinant *R. vannielii* EF-P production, *efp* was cloned into the pET expression plasmid and overproduced using the pET expression system in *E. coli* BL21 (DE3) (Invitrogen). Cells were harvested after cultivation in LB supplemented with IPTG (1 mM) at 18°C overnight, and the resulting pellet was frozen at −80 °C. Cells with chromosomally incorporated His-tagged *efp* (*R. vannielii* EF-P and *W. virosa* EF-P) were grown in LB until the mid-exponential growth phase, harvested and kept at −80 °C until further processing.

For lysis, cells were resuspended in 0.1 M, pH 7.6 sodium phosphate buffer (supplemented with 300 mM NaCl and DNase) and lysed using the high-pressure cell disrupter (Constant Systems), by running the sample twice under 1.9 kbar with subsequent cell fractionation using ultracentrifugation. The His-tagged proteins were purified using the QIAexpress Ni-NTA Protein Purification System (Qiagen) with 0.1 M, pH 7.6 sodium phosphate buffer supplemented with 300 mM NaCl and 30 mM - 200 mM imidazole. Samples were dialyzed to remove imidazole at 4°C for 24 h using SERVAPORE dialysis tubing (MWCO 12000-14000, SERVA) according to manufacturer’s instructions. Proteins were concentrated using the Amicon^®^ Ultra-15 Centrifugal Filter Units, 3kDa (Millipore).

### SDS-PAGE and Western Blot analysis

Proteins were separated by sodium dodecyl sulfate-polyacrylamide gel electrophoresis (SDS-PAGE) ^48^ using 12.5 % Bis-tris acrylamide gels. Proteins were transferred to a nitrocellulose membrane using the wet-blot apparatus (Mini Trans-Blot Cell, Bio-Rad). *E. c.* EF-P was detected by incubating the membrane in Tris-buffered saline (TBS) with primary polyclonal antibodies against *E. coli* EF-P (rabbit, Eurogentec) with final concentration of 1:5,000, whereas His-tagged EF-P was detected with primary monoclonal antibodies against the His-Tag (1:10,000; AB_2536841, Invitrogen). Membranes were incubated with the secondary antibodies conjugated with a fluorophore (1:20,000 in TBS) (anti mouse: ab216776; anti rabbit: ab216773) and imaged using the Odyssey CLx (LI-COR Biosciences). For detection of *R. vannielii* EF-P, recombinantly produced EF-P was purified and sent to Eurogentec for polyclonal antibody generation (Speedy 28-day program in rabbits, Eurogentec). The blood serum containing antibodies against the *R. vannielii* EF-P (α 652) was diluted to final concentration of 1:100.

### Mass spectrometry for identification of modification status

For top-down EF-P measurements the purified proteins were desalted on the ZipTip with C4 resin (Millipore, ZTC04S096) and eluted with 50 % (v/v) acetonitrile 0.1 % (v/v) formic acid (FA) buffer resulting in ∼10 μM final protein concentration in 200–400 μl total volume. MS measurements were performed on an Orbitrap Eclipse Tribrid Mass Spectrometer (Thermo Fisher Scientific) via direct injection, a HESI-Spray source (Thermo Fisher Scientific) and FAIMS interface (Thermo Fisher Scientific) in a positive, peptide mode. Typically, the FAIMS compensation voltage (CV) was optimized by a continuous scan. The most intense signal was usually obtained at −5 CV. The MS spectra were acquired with at least 120,000 FWHM, AGC target 100 and 2-5 microscans covering the 800-1100 m/z range. The spectra were deconvoluted in Freestyle (Thermo) using the Xtract Deconvolution algorithm.

For bottom-up proteomics, samples were prepared in 96-well plate using the optimized SP3 protocol ^49^. The purified EF-P sample (1 μL, 10 μM) was diluted to total volume of 50 μL with 1 % NP40, 0.2 % (w/v) SDS in 25 mM HEPES, pH 7.5. The protein was loaded onto a mixture of hydrophilic and hydrophobic carboxylate-coated magnetic beads (10 μL each) pre-washed three times with 100 µL of MS-grade H_2_O. The magnetic beads with protein sample were mixed at 850 rpm, 1 min at room temperature (RT). To initiate the binding, 60 µL of absolute EtOH was added, and the mixture was incubated at RT for 5 min at 850 rpm. Subsequently, the beads were washed three times with 80 % (v/v) EtOH, with incubation at RT for 1 min and 850 rpm between each wash. After the last wash the beads were resuspended in 50 μL of 100 mM ammonium acetate buffer (ABC). The proteins were reduced and alkylated by addition of 5 μL of 100 mM tris(2-carboxyethyl) phosphine (TCEP) and 5 μL of 400 mM chloroacetamide (CAA) and incubation at 95 °C for 5 min, 850 rpm. Samples were cooled to RT. The on-beads digestion was performed with either trypsin or chymotrypsin. Chymotrypsin digestion: ABC buffer was supplemented with 10 mM CaCl_2_, 1 μg of chymotrypsin (Thermo Scientific, 90056) and incubated at 25 °C overnight. The resulting peptide mixture was eluted from the magnetic beads into a new 1.5 mL tube. The magnetic beads were washed with 50 and 30 μL of 1 % (v/v) formic acid and incubated at 40 °C, 850 rpm for 5 min. The fractions were added to the first elution fraction. The combined fractions were further purified from remaining magnetic beads. Alternatively, for *R. v.* EF-P overproduced in *E. coli*, the EF-P was directly digested in ABC buffer (50 μL) supplemented with 10 mM CaCl_2_ and 1 μg of chymotrypsin (Thermo Scientific, 90056), without previous clean-up on magnetic beads.

MS measurements were performed on an Orbitrap Eclipse Tribrid Mass Spectrometer (Thermo Fisher Scientific) coupled to an UltiMate 3000 Nano-HPLC (Thermo Fisher Scientific) via a nanospray Flex ion source (Thermo Fisher Scientific) equipped with column oven (Sonation) and FAIMS interface (Thermo Fisher Scientific). Peptides were loaded on an Acclaim PepMap 100 μ-precolumn cartridge (5 μm, 100 Å, 300 μm ID x 5 mm, Thermo Fisher Scientific) and separated at 40°C on a PicoTip emitter (noncoated, 15 cm, 75 μm ID, 8 μm tip, New Objective) that was *in-house* packed with Reprosil-Pur 120 C18-AQ material (1.9 μm, 150 Å, Dr. A. Maisch GmbH). Buffer composition. Buffer A consists of MS-grade H_2_O supplemented with 0.1 % FA. Buffer B consists of acetonitrile supplemented with 0.1 % FA. The 41-or 126-min LC gradient from 4 to 35.2 % buffer B was used. The flow rate was 0.3 µL/min.

#### Data-independent acquisition

The DIA duty cycle consisted of one MS1 scan followed by 30 MS2 scans with an isolation window of the 4 m/z range, overlapping with an adjacent window at the 2 m/z range. MS1 scan was conducted with Orbitrap at 60000 resolution power and a scan range of 200 – 1800 m/z with an adjusted RF lens at 30 %. MS2 scans were conducted with Orbitrap at 30000 resolution power, RF lens was set to 30 %. The precursor mass window was restricted to a 500 – 740 m/z range. HCD fragmentation was enabled as an activation method with a fixed collision energy of 35 %. FAIMS was performed with one CV at −45V for both MS1 and MS2 scans during the duty cycle.

#### Data-dependent acquisition

(*R. v.* EF-P overproduced in *E. coli*) For measurements of DDA, the Orbitrap Eclipse Tribrid Mass Spectrometer was operated with the following settings: Polarity: positive; MS1 resolution: 240k; MS1 AGC target: standard; MS1 maximum injection time: 50 ms; MS1 scan range: m/z 375-1500; MS2 ion trap scan rate: rapid; MS2 AGC target: standard; MS2 maximum injection time: 35 ms; MS2 cycle time: 1.7 s; MS2 isolation window: m/z 1.2; HCD stepped normalised collision energy: 30 %; intensity threshold: 1.0e4 counts; included charge states: 2-6; dynamic exclusion: 60 s. FAIMS was performed with two alternating CVs, including - 50 V and −70 V.

##### Mass spectrometry for identification of modification status

###### Computational evaluation of DIA raw files

Raw files were converted in the first step with “MSConvertGUI” as a part of the “ProteoWizard” software package (http://www.proteowizard.org/download.html) to an output mzML format applying the “peakPicking” filter with “vendor msLevel=1”, and the “Demultiplex” filter with parameters “Overlap Only” and “mass error” set to 10 ppm.

###### Standalone DIA-NN software under version 1.8.1 was used for protein identification and quantification

First, a spectral library was predicted *in silico* by the software’s deep learning-based spectra, RTs and IMs prediction using Uniprot *E. coli* decoyed FASTA (canonical and isoforms with added *R. vannielii* EF-P sequence). DIA-NN search settings: FASTA digest for library-free search/library generation option was enabled, together with a match between runs (MBR) option and precursor FDR level set at 1 %. Library generation was set to smart profiling, Quantification strategy - Robust LC. The mass accuracy and the scan window were set to 0 to allow the software to identify optimal conditions. The precursor m/z range was changed to 500-740 m/z to fit the measuring parameters. Carbamidomethylation was set as a fixed modification, oxidation of methionine and N-term acetylation were set as variable modifications. On the contrary, the small-scale samples of the 96-well plate were calculated without carbamidomethylation as a fixed modification.

###### Computational evaluation of DDA raw files

MS Raw files were analyzed using MaxQuant software. Searches were performed against the Uniprot database for *E. coli* (*R. vannielii* EF-P sequence). At least two unique peptides were required for protein identification. False discovery rate determination was carried out using a decoy database and thresholds were set to 1 % FDR both at peptide-spectrum match and at protein levels.

### Mass spectrometry for proteome analysis

Cells were cultivated in LB under constant shaking at 37 °C until reaching the exponential growth phase. Cells were harvested (OD_600_=0.5) and were processed with the iST kit (Preomics) as recommended by the manufacturer. Samples were evaporated to dryness, resuspended in LC-LOAD buffer to 0.2µg/µL and injected in an Ultimate 3000 RSLCnano system (Thermo) separated in a 25-cm Aurora column (Ionopticks) with a 100-min gradient from 4 to 40 % acetonitrile in 0.1 % formic acid. The effluent from the HPLC was directly electrosprayed into an Orbitrap Exploris 480 (Thermo) operated in data dependent mode to automatically switch between full scan MS and MS/MS acquisition. Survey full scan MS spectra (from m/z 350-1200) were acquired with a resolution of R=60,000 at m/z 400 (AGC target of 3×10^6^). The 20 most intense peptide ions with charge states between 2 and 6 were sequentially isolated to a target value of 1×10^5^ and fragmented at 30 % normalized collision energy. Typical mass spectrometric conditions were: spray voltage, 1.5 kV; no sheath and auxiliary gas flow; heated capillary temperature, 275°C; intensity selection threshold, 3×10^5^.

The MaxQuant 2.1.0.0 software was used for protein identification and quantification by label-free quantification (LFQ)^29^ with the following parameters: Database Uniprot_UP000000625_Ecoli_20220309.fasta including the *Rhodomicrobium vannielii* EFP sequence; MS tol, 10 ppm; MS/MS tol, 20 ppm Da; Peptide FDR, 0.1; Protein FDR, 0.01 min; Peptide Length, 7; Variable modifications, Oxidation (M); Fixed modifications, Carbamidomethyl (C); Peptides for protein quantitation, razor and unique; Min. peptides, 1; Min. ratio count, 2. For display and analysis, the Perseus software ^50,51^ was used. A list as a reference for proteins with polyproline-motifs in *E. coli* was taken from Qi *et* al. ^13^. Data have been uploaded to the PRIDE repository ^52^ <u>PXD044929.</u>

### Statistical analysis

All measurements, except intact protein mass measurements and chymotrypsin digestions, are from at least three biological replicates. Statistical analysis was calculated using Microsoft Office *Excel* 2019 (student’s unpaired two-sided t-test; 95 % confidence intervals). Values were considered as significantly different when the calculated *p-value* was below 0.05.

Perseus (2.0.9.0) was used to log_2_ transform LFQ intensities, replace missing values from normal distribution and construct the volcano plots. To determine which proteins were differentially expressed between experimental conditions, we applied a t-test with a permutation-based FDR calculation with n=3, FDR=0.05 and s0=0.1 to the log 2 LFQ protein values, where s0 controls the relative importance of t-test p-value and difference between means. At s0=0 only the p-value matters, while at nonzero s0 also the difference of means plays a role ^53^.

## Data availability

The mass spectrometry proteomics data have been deposited to the ProteomeXchange Consortium via the PRIDE ^52^ partner repository with the dataset identifier PXD044929.

## Acknowledgements

We thank Prof. Bernhard Schink from University of Konstanz for the support in handling *R. vannielii*. We also thank Haoyu Chen and Thomas Krause for excellent technical assistance. This work was financially supported by Deutsche Forschungsgemeinschaft (DFG, German Research Foundation) grant JU270/20-1, project number 449926427, RTG2062, (Molecular Principles of Synthetic Biology) to K.J. and SFB 1309, project number 325871075 to K.J., P.K. and A.I.

## Author Contributions

Conceptualization, U.T. and K.J.; Methodology, U.T., P.K., I.F., A.I., K.J.; Investigation, U.T., P.K., R.K., I.F.; Writing – Original Draft, U.T., K.J.; Funding acquisition and Resources, P.K., A.I., K.J.; Supervision, K.J.

## References

1. Hanson, G., and Coller, J. (2018). Codon optimality, bias and usage in translation and mRNA decay. Nat Rev Mol Cell Biol 19, 20–30. 10.1038/nrm.2017.91.

2. Doerfel, L.K., Wohlgemuth, I., Kubyshkin, V., Starosta, A.L., Wilson, D.N., Budisa, N., and Rodnina, M.V. (2015). Entropic Contribution of Elongation Factor P to Proline Positioning at the Catalytic Center of the Ribosome. J Am Chem Soc 137, 12997–13006. 10.1021/jacs.5b07427.

3. Muto, H., and Ito, K. (2008). Peptidyl-prolyl-tRNA at the ribosomal P-site reacts poorly with puromycin. Biochem Biophys Res Commun 366, 1043–1047. 10.1016/j.bbrc.2007.12.072.

4. Pavlov, M.Y., Watts, R.E., Tan, Z., Cornish, V.W., Ehrenberg, M., and Forster, A.C. (2009). Slow peptide bond formation by proline and other N-alkylamino acids in translation. Proc Natl Acad Sci U S A 106, 50–54. 10.1073/pnas.0809211106.

5. Wohlgemuth, I., Brenner, S., Beringer, M., and Rodnina, M.V. (2008). Modulation of the rate of peptidyl transfer on the ribosome by the nature of substrates. J Biol Chem 283, 32229–32235. 10.1074/jbc.M805316200.

6. Doerfel, L.K., Wohlgemuth, I., Kothe, C., Peske, F., Urlaub, H., and Rodnina, M.V. (2013). EF-P Is Essential for Rapid Synthesis of Proteins Containing Consecutive Proline Residues. Science 339, 85–88. 10.1126/science.1229017.

7. Ude, S., Lassak, J., Starosta, A.L., Kraxenberger, T., Wilson, D.N., and Jung, K. (2013). Translation elongation factor EF-P alleviates ribosome stalling at polyproline stretches. Science 339, 82–85. 10.1126/science.1228985.

8. Tanner, D.R., Cariello, D.A., Woolstenhulme, C.J., Broadbent, M.A., and Buskirk, A.R. (2009). Genetic Identification of Nascent Peptides That Induce Ribosome Stalling. Journal of Biological Chemistry 284, 34809–34818. 10.1074/jbc.M109.039040.

9. Peil, L., Starosta, A.L., Lassak, J., Atkinson, G.C., Virumae, K., Spitzer, M., Tenson, T., Jung, K., Remme, J., and Wilson, D.N. (2013). Distinct XPPX sequence motifs induce ribosome stalling, which is rescued by the translation elongation factor EF-P. Proc Natl Acad Sci U S A 110, 15265–15270. 10.1073/pnas.1310642110.

10. Adzhubei, A.A., Sternberg, M.J., and Makarov, A.A. (2013). Polyproline-II helix in proteins: structure and function. J Mol Biol 425, 2100–2132. 10.1016/j.jmb.2013.03.018.

11. Starosta, A.L., Lassak, J., Peil, L., Atkinson, G.C., Woolstenhulme, C.J., Virumae, K., Buskirk, A., Tenson, T., Remme, J., Jung, K., and Wilson, D.N. (2014). A conserved proline triplet in Val-tRNA synthetase and the origin of elongation factor P. Cell Rep 9, 476–483. 10.1016/j.celrep.2014.09.008.

12. Motz, M., and Jung, K. (2018). The role of polyproline motifs in the histidine kinase EnvZ. PLoS One 13, e0199782. 10.1371/journal.pone.0199782.

13. Qi, F., Motz, M., Jung, K., Lassak, J., and Frishman, D. (2018). Evolutionary analysis of polyproline motifs in *Escherichia coli* reveals their regulatory role in translation. PLoS Comput Biol 14, e1005987. 10.1371/journal.pcbi.1005987.

14. Gutierrez, E., Shin, B.S., Woolstenhulme, C.J., Kim, J.R., Saini, P., Buskirk, A.R., and Dever, T.E. (2013). eIF5A promotes translation of polyproline motifs. Mol Cell 51, 35–45. 10.1016/j.molcel.2013.04.021.

15. Huter, P., Arenz, S., Bock, L.V., Graf, M., Frister, J.O., Heuer, A., Peil, L., Starosta, A.L., Wohlgemuth, I., Peske, F., et al. (2017). Structural Basis for Polyproline-Mediated Ribosome Stalling and Rescue by the Translation Elongation Factor EF-P. Molecular Cell 68, 515-+. 10.1016/j.molcel.2017.10.014.

16. Blaha, G., Stanley, R.E., and Steitz, T.A. (2009). Formation of the first peptide bond: the structure of EF-P bound to the 70S ribosome. Science 325, 966–970. 10.1126/science.1175800.

17. Park, M.H., Cooper, H.L., and Folk, J.E. (1982). The biosynthesis of protein-bound hypusine (N epsilon −(4-amino-2-hydroxybutyl)lysine). Lysine as the amino acid precursor and the intermediate role of deoxyhypusine (N epsilon −(4-aminobutyl)lysine). J Biol Chem 257, 7217–7222.

18. Behshad, E., Ruzicka, F.J., Mansoorabadi, S.O., Chen, D., Reed, G.H., and Frey, P.A. (2006). Enantiomeric free radicals and enzymatic control of stereochemistry in a radical mechanism: the case of lysine 2,3-aminomutases. Biochemistry 45, 12639–12646. 10.1021/bi061328t.

19. Yanagisawa, T., Sumida, T., Ishii, R., Takemoto, C., and Yokoyama, S. (2010). A paralog of lysyl-tRNA synthetase aminoacylates a conserved lysine residue in translation elongation factor P. Nat Struct Mol Biol 17, 1136–1143. 10.1038/nsmb.1889.

20. Navarre, W.W., Zou, S.B., Roy, H., Xie, J.L., Savchenko, A., Singer, A., Edvokimova, E., Prost, L.R., Kumar, R., Ibba, M., and Fang, F.C. (2010). PoxA, yjeK, and elongation factor P coordinately modulate virulence and drug resistance in *Salmonella enterica*. Mol Cell 39, 209–221. 10.1016/j.molcel.2010.06.021.

21. Peil, L., Starosta, A.L., Virumae, K., Atkinson, G.C., Tenson, T., Remme, J., and Wilson, D.N. (2012). Lys34 of translation elongation factor EF-P is hydroxylated by YfcM. Nat Chem Biol 8, 695–697. 10.1038/nchembio.1001.

22. Roy, H., Zou, S.B., Bullwinkle, T.J., Wolfe, B.S., Gilreath, M.S., Forsyth, C.J., Navarre, W.W., and Ibba, M. (2011). The tRNA synthetase paralog PoxA modifies elongation factor-P with (R)-beta-lysine. Nat Chem Biol 7, 667–669. 10.1038/nchembio.632.

23. Lassak, J., Keilhauer, E.C., Furst, M., Wuichet, K., Godeke, J., Starosta, A.L., Chen, J.M., Sogaard-Andersen, L., Rohr, J., Wilson, D.N., et al. (2015). Arginine-rhamnosylation as new strategy to activate translation elongation factor P. Nat Chem Biol 11, 266–270. 10.1038/nchembio.1751.

24. Rajkovic, A., Erickson, S., Witzky, A., Branson, O.E., Seo, J., Gafken, P.R., Frietas, M.A., Whitelegge, J.P., Faull, K.F., Navarre, W., et al. (2015). Cyclic Rhamnosylated Elongation Factor P Establishes Antibiotic Resistance in *Pseudomonas aeruginosa*. mBio 6, e00823. 10.1128/mBio.00823-15.

25. Rajkovic, A., Hummels, K.R., Witzky, A., Erickson, S., Gafken, P.R., Whitelegge, J.P., Faull, K.F., Kearns, D.B., and Ibba, M. (2016). Translation Control of Swarming Proficiency in *Bacillus subtilis* by 5-Amino-pentanolylated Elongation Factor P. J Biol Chem 291, 10976–10985. 10.1074/jbc.M115.712091.

26. Hummels, K.R., Witzky, A., Rajkovic, A., Tollerson, R., 2nd, Jones, L.A., Ibba, M., and Kearns, D.B. (2017). Carbonyl reduction by YmfI in *Bacillus subtilis* prevents accumulation of an inhibitory EF-P modification state. Mol Microbiol 106, 236–251. 10.1111/mmi.13760.

27. Pinheiro, B., Scheidler, C.M., Kielkowski, P., Schmid, M., Forne, I., Ye, S., Reiling, N., Takano, E., Imhof, A., Sieber, S.A., et al. (2020). Structure and Function of an Elongation Factor P Subfamily in Actinobacteria. Cell Rep 30, 4332–4342 e4335. 10.1016/j.celrep.2020.03.009.

28. Tollerson, R., 2nd, Witzky, A., and Ibba, M. (2018). Elongation factor P is required to maintain proteome homeostasis at high growth rate. Proc Natl Acad Sci U S A 115, 11072–11077. 10.1073/pnas.1812025115.

29. Cox, J., Hein, M.Y., Luber, C.A., Paron, I., Nagaraj, N., and Mann, M. (2014). Accurate proteome-wide label-free quantification by delayed normalization and maximal peptide ratio extraction, termed MaxLFQ. Mol Cell Proteomics 13, 2513–2526. 10.1074/mcp.M113.031591.

30. Volkwein, W., Krafczyk, R., Jagtap, P.K.A., Parr, M., Mankina, E., Macosek, J., Guo, Z., Furst, M., Pfab, M., Frishman, D., et al. (2019). Switching the Post-translational Modification of Translation Elongation Factor EF-P. Front Microbiol 10, 1148. 10.3389/fmicb.2019.01148.

31. Hanawa-Suetsugu, K., Sekine, S., Sakai, H., Hori-Takemoto, C., Terada, T., Unzai, S., Tame, J.R., Kuramitsu, S., Shirouzu, M., and Yokoyama, S. (2004). Crystal structure of elongation factor P from *Thermus thermophilus* HB8. Proc Natl Acad Sci U S A 101, 9595–9600. 10.1073/pnas.0308667101.

32. Choi, S., and Choe, J. (2011). Crystal structure of elongation factor P from *Pseudomonas aeruginosa* at 1.75 A resolution. Proteins 79, 1688–1693. 10.1002/prot.22992.

33. Jumper, J., Evans, R., Pritzel, A., Green, T., Figurnov, M., Ronneberger, O., Tunyasuvunakool, K., Bates, R., Zidek, A., Potapenko, A., et al. (2021). Highly accurate protein structure prediction with AlphaFold. Nature 596, 583-+. 10.1038/s41586-021-03819-2.

34. Varadi, M., Anyango, S., Deshpande, M., Nair, S., Natassia, C., Yordanova, G., Yuan, D., Stroe, O., Wood, G., Laydon, A., et al. (2022). AlphaFold Protein Structure Database: massively expanding the structural coverage of protein-sequence space with high-accuracy models. Nucleic Acids Res 50, D439–D444. 10.1093/nar/gkab1061.

35. UniProt, C. (2023). UniProt: the Universal Protein Knowledgebase in 2023. Nucleic Acids Res 51, D523–D531. 10.1093/nar/gkac1052.

36. Hayter, J.R., Robertson, D.H., Gaskell, S.J., and Beynon, R.J. (2003). Proteome analysis of intact proteins in complex mixtures. Mol Cell Proteomics 2, 85–95. 10.1074/mcp.M200078-MCP200.

37. Lee, S.W., Berger, S.J., Martinovic, S., Pasa-Tolic, L., Anderson, G.A., Shen, Y., Zhao, R., and Smith, R.D. (2002). Direct mass spectrometric analysis of intact proteins of the yeast large ribosomal subunit using capillary LC/FTICR. Proc Natl Acad Sci U S A 99, 5942–5947. 10.1073/pnas.082119899.

38. Pfab, M., Kielkowski, P., Krafczyk, R., Volkwein, W., Sieber, S.A., Lassak, J., and Jung, K. (2021). Synthetic post-translational modifications of elongation factor P using the ligase EpmA. FEBS J 288, 663–677. 10.1111/febs.15346.

39. Reiling, N., Homolka, S., Walter, K., Brandenburg, J., Niwinski, L., Ernst, M., Herzmann, C., Lange, C., Diel, R., Ehlers, S., and Niemann, S. (2013). Clade-specific virulence patterns of *Mycobacterium tuberculosis* complex strains in human primary macrophages and aerogenically infected mice. mBio 4. 10.1128/mBio.00250-13.

40. Bradshaw, E., Saalbach, G., and McArthur, M. (2013). Proteomic survey of the *Streptomyces coelicolor* nucleoid. J Proteomics 83, 37–46. 10.1016/j.jprot.2013.02.033.

41. Wards, B.J., and Collins, D.M. (1996). Electroporation at elevated temperatures substantially improves transformation efficiency of slow-growing mycobacteria. FEMS Microbiol Lett 145, 101–105. 10.1111/j.1574-6968.1996.tb08563.x.

42. Katz, A., Solden, L., Zou, S.B., Navarre, W.W., and Ibba, M. (2014). Molecular evolution of protein-RNA mimicry as a mechanism for translational control. Nucleic Acids Res 42, 3261–3271. 10.1093/nar/gkt1296.

43. Marman, H.E., Mey, A.R., and Payne, S.M. (2014). Elongation factor P and modifying enzyme PoxA are necessary for virulence of *Shigella flexneri*. Infect Immun 82, 3612–3621. 10.1128/IAI.01532-13.

44. Guo, Q., Cui, B., Wang, M., Li, X., Tan, H., Song, S., Zhou, J., Zhang, L.H., and Deng, Y. (2022). Elongation factor P modulates *Acinetobacter baumannii* physiology and virulence as a cyclic dimeric guanosine monophosphate effector. Proc Natl Acad Sci U S A 119, e2209838119. 10.1073/pnas.2209838119.

45. Lassak, J., Henche, A.L., Binnenkade, L., and Thormann, K.M. (2010). ArcS, the cognate sensor kinase in an atypical Arc system of *Shewanella oneidensis* MR-1. Appl Environ Microbiol 76, 3263–3274. 10.1128/AEM.00512-10.

46. Keseler, I.M., Gama-Castro, S., Mackie, A., Billington, R., Bonavides-Martinez, C., Caspi, R., Kothari, A., Krummenacker, M., Midford, P.E., Muniz-Rascado, L., et al. (2021). The EcoCyc Database in 2021. Front Microbiol 12, 711077. 10.3389/fmicb.2021.711077.

47. Miller, J.H. (1992). A short course in bacterial genetics : a laboratory manual and handbook for Escherichia coli and related bacteria (Cold Spring Harbor Laboratory Press).

48. Laemmli, U.K. (1970). Cleavage of structural proteins during the assembly of the head of bacteriophage T4. Nature 227, 680–685. 10.1038/227680a0.

49. Hughes, C.S., Moggridge, S., Muller, T., Sorensen, P.H., Morin, G.B., and Krijgsveld, J. (2019). Single-pot, solid-phase-enhanced sample preparation for proteomics experiments. Nat Protoc 14, 68–85. 10.1038/s41596-018-0082-x.

50. Tyanova, S., Temu, T., Sinitcyn, P., Carlson, A., Hein, M.Y., Geiger, T., Mann, M., and Cox, J. (2016). The Perseus computational platform for comprehensive analysis of (prote)omics data. Nat Methods 13, 731–740. 10.1038/nmeth.3901.

51. Tyanova, S., and Cox, J. (2018). Perseus: A Bioinformatics Platform for Integrative Analysis of Proteomics Data in Cancer Research. Methods Mol Biol 1711, 133–148. 10.1007/978-1-4939-7493-1_7.

52. Perez-Riverol, Y., Bai, J., Bandla, C., Garcia-Seisdedos, D., Hewapathirana, S., Kamatchinathan, S., Kundu, D.J., Prakash, A., Frericks-Zipper, A., Eisenacher, M., et al. (2022). The PRIDE database resources in 2022: a hub for mass spectrometry-based proteomics evidences. Nucleic Acids Res 50, D543–D552. 10.1093/nar/gkab1038.

53. Tusher, V.G., Tibshirani, R., and Chu, G. (2001). Significance analysis of microarrays applied to the ionizing radiation response. Proc Natl Acad Sci U S A 98, 5116–5121. 10.1073/pnas.091062498.

