## Supplemental Information for "Decrypting the functional design of unmodified translation elongation factor P"

**Supplementary Table S1. Protein sequence identity of *E. coli* EF-P and EF-Ps of the PGKGP-subfamily.** Multiple sequence alignment and percent identity matrix were calculated using the Multiple Sequence Alignment Tool from Clustal Omega (Clustal 2.1) and is given in percentage (%).

| | ***E. c.*** | ***C. h.*** | ***C. l.*** | ***C. a.*** | ***C. w.*** | ***P. g.*** | ***R. v.*** | ***S. v.*** | ***W. v.*** |
|---|---|---|---|---|---|---|---|---|---|
| ***E. c.*** | | 44.92 | 44.15 | 39.13 | 40.54 | 41.08 | 32.09 | 41.94 | 37.84 |
| ***C. h.*** | 44.92 | | 88.83 | 40.86 | 44.86 | 43.01 | 40.64 | 43.32 | 40.86 |
| ***C. l.*** | 44.15 | 88.83 | | 40.86 | 45.95 | 40.86 | 37.97 | 42.25 | 40.32 |
| ***C. a.*** | 39.13 | 40.86 | 40.86 | | 44.86 | 53.19 | 32.97 | 43.01 | 66.49 |
| ***C. w.*** | 40.54 | 44.86 | 45.95 | 44.86 | | 45.65 | 35.14 | 46.49 | 50.54 |
| ***P. g.*** | 41.08 | 43.01 | 40.86 | 53.19 | 45.65 | | 35.68 | 49.46 | 57.98 |
| ***R. v.*** | 32.09 | 40.64 | 37.97 | 32.97 | 35.14 | 35.68 | | 36.56 | 31.89 |
| ***S. v.*** | 41.94 | 43.32 | 42.25 | 43.01 | 46.49 | 49.46 | 36.56 | | 41.94 |
| ***W. v.*** | 37.84 | 40.86 | 40.32 | 66.49 | 50.54 | 57.98 | 31.89 | 41.94 | |

**E. c.** - *Escherichia coli*
**C. h.** - *Campylobacter hominis*
**C. l.** - *Campylobacter lari*
**C. a.** - *Cellulophaga algicola*
**C. w.** - *Conexibacter woesei*
**P. g.** - *Porphyromonas gingivalis*
**R. v.** - *Rhodomicrobium vannielii*
**S. v.** - *Streptomyces venezuelae*
**W. v.** - *Weeksella virosa*

**Supplementary Table S2. Doubling times of *E. coli* wild type (wt) and mutants.** Strains were grown as described in **Fig. 2c**.

| | **Doubling time (min)** |
|---|---|
| wt | 24.67 |
| Δ*efp*::*R. v. efp* | 25.82 |
| Δ*efp*::*R. v. efp* Δ*epmA* | 25.86 |
| Δ*epmA* | 32.90 |

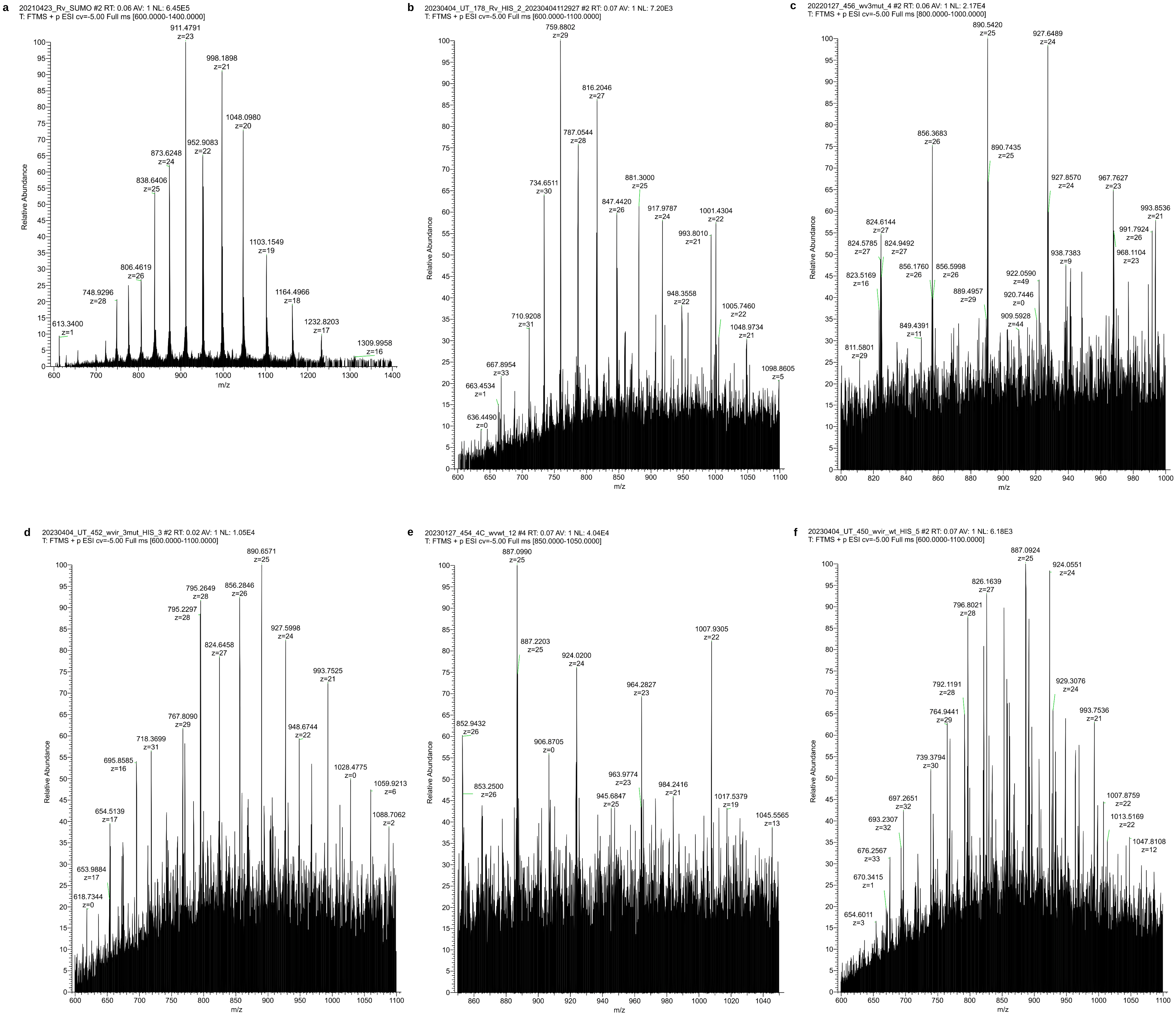


**Supplementary Figure S1. Mass spectrometry analysis of EF-Ps produced in *E. coli*.** **a** Intact mass spectra of recombinantly overproduced *R. vannielii* EF-P purified from *E. coli* wt. **b** Intact mass spectra of His-tagged *R. vannielii* EF-P (chromosomally encoded), purified from *E. coli* wt. **c** Intact mass spectra of His-tagged *W. virosa* EF-P variant with substitutions S1-S3 (chromosomally encoded), purified from *E. coli* Δ*epmA*. **d** Intact mass spectra of His-tagged *W. virosa* EF-P variant with substitutions S1-S3 (chromosomally encoded), purified from *E. coli* wt. **e** Intact mass spectra of His-tagged *W. virosa* EF-P wt (chromosomally encoded), purified from *E. coli* Δ*epmA*. **f** Intact mass spectra of His-tagged *W. virosa* EF-P wt (chromosomally encoded), purified from *E. coli* wt. S – substitution (S1 – P34Q; S2 – V49R; S3 – E62K).

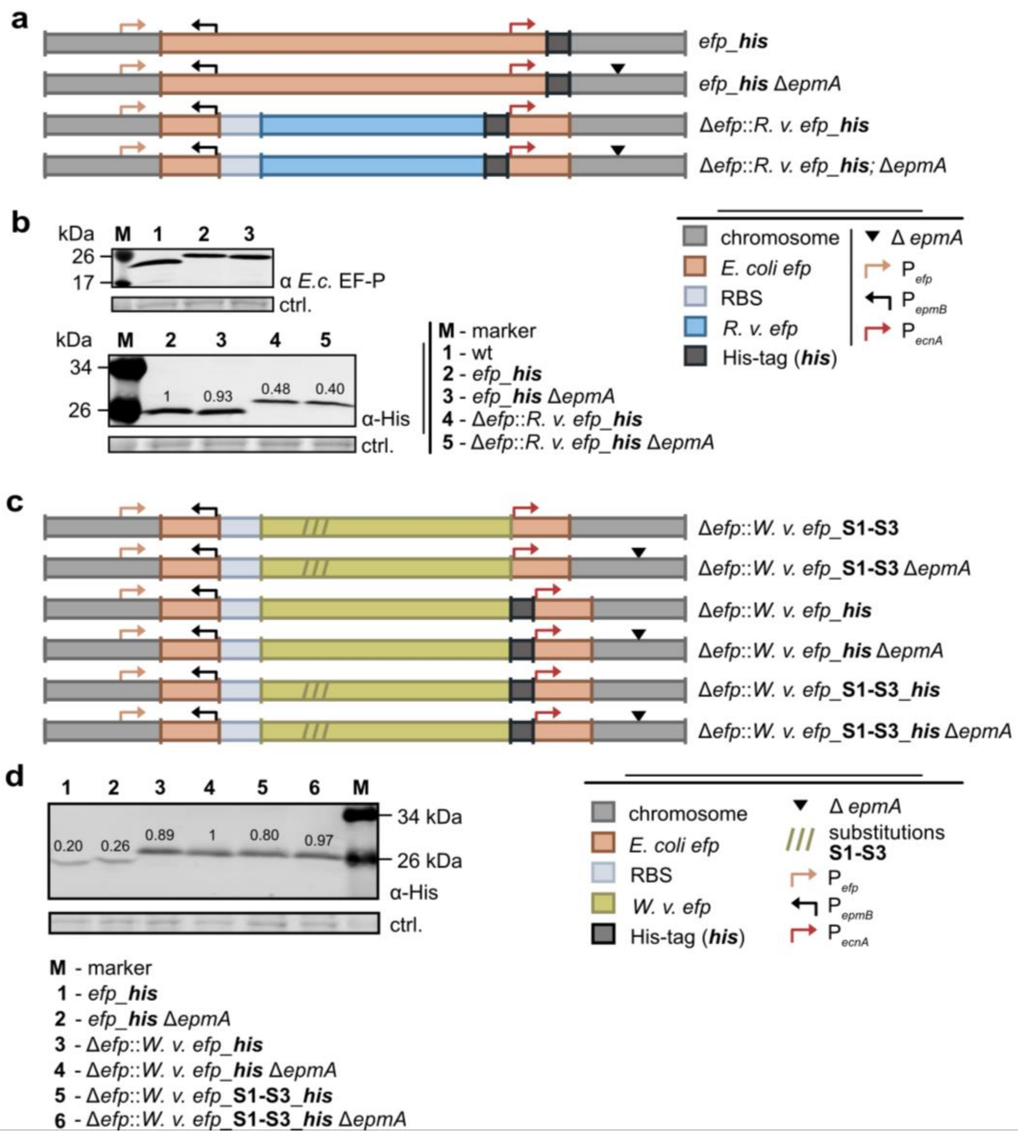


**Supplementary Figure S2. Protein production of His-tagged EF-P variants in *E. coli*. a** Schematic overview of the constructed *E. coli* mutants expressing *his*-tagged *efp* or *R. v. efp*. **b** Western blot analysis of *R. v.* EF-P production in *E. coli* mutants after growth in LB to mid-exponential phase using antibodies against *E. coli* EF-P (upper graph) or the His-tag (lower graph).**c** Schematic overview of the constructed *E. coli* mutants expressing *his*-tagged *efp* variants of *W. v.* **d** Western blot analysis of *W. v.* EF-P production in *E. coli* mutants after growth in LB to mid-exponential phase using antibodies against the His-tag. Relative band intensities calculated using ImageJ are displayed above the corresponding protein bands. Protein bands corresponding to a 72 kDa protein after staining with 2,2,2-trichloroethanol (TCE) were used as loading controls (ctrl.). Gene insertions in **a** and **c** are depicted in colored boxes, and promoter locations as coloured arrows. S1-S3 - amino acid substitutions (S1 – P34Q; S2 – V49R; S3 – E62K).

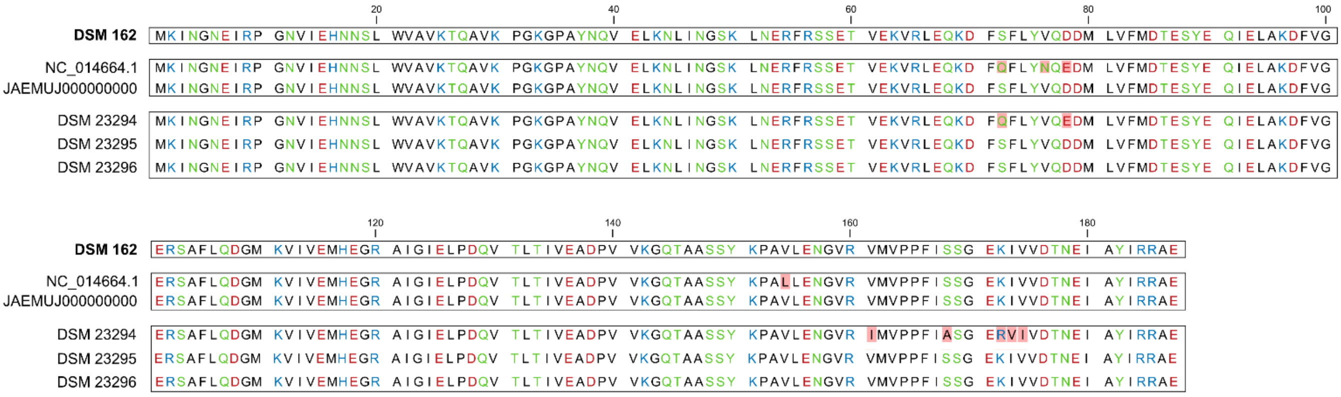


**Supplementary Figure S3. Comparison of the sequence of EF-Ps from different *R. vannielii* strains and the publicly available sequences**. Different colors represent the polarity of the amino acids (black-hydrophobic, light green-hydrophilic, red-acidic, blue-basic). Differences in amino acids between strains are highlighted in pink. Discrepancies between the EF-P sequences of the type strain DSM 162 used in this study and the publicly available sequence of *R. vannielii* ATCC171000 (GenBank accession number NC_014664.1) were ruled out by comparison with the revised EF-P sequence of *R. vannielii* ATCC171000 (GenBank accession number JAEMUJ000000000.1 [1]), and the sequencing results of EF-P from evolutionarily related strains (DSM 23294, DSM 23295, and DSM 23296).

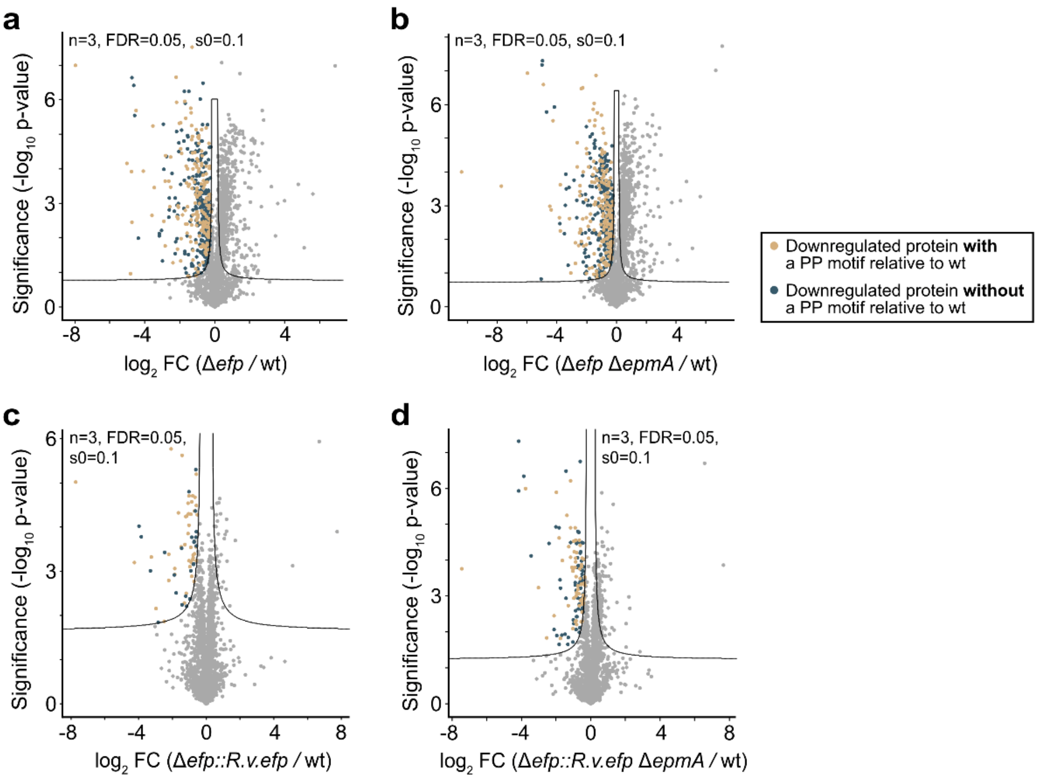


**Supplementary Figure S4. Proteome-wide analysis of polyproline-containing proteins in the *E. coli* mutants Δ*efp*, Δ*efp* Δ*epmA* after complementation with *R. v. efp*. a-d** Volcano plot analysis highlight the downregulation of proteins with/without polyproline motifs (PP-motifs) in *E. coli* mutants Δ*efp* (**a**), Δ*efp* Δ*epmA* (**b**), Δ*efp*::*R. v. efp* (**c**) and Δ*efp*::*R. v. efp* Δ*epmA* (**d**) compared to wild type (wt). The x-axes show for each protein the fold change (FC) of the mean value of the $log_2$ protein intensity (LFQ) between two strains. The y-axes show the significance level of the observed difference between the two strains (-$log_{10}$ p-value of the t-test). The test was adjusted for multiple comparisons (permutation-based FDR with n=3, FDR=0.05 and s0=0.1).

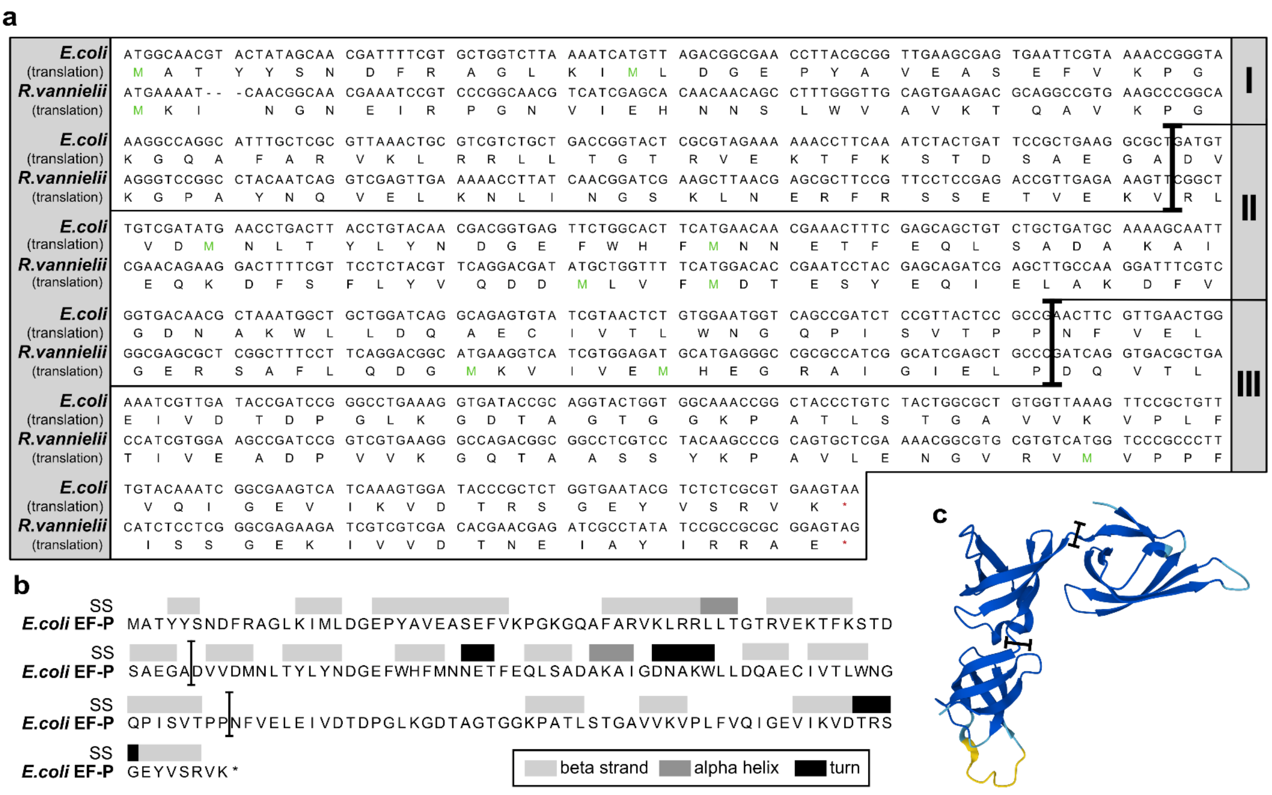


**Supplementary Figure S5. Construction principles of *R. vannielii* EF-P and *E. coli* EF-P hybrids.** **a** Nucleotide and protein sequence alignment of *E. coli* EF-P and *R. vannielii* EF-P with selected boundaries (black bulk arrows) for each EF-P domain (I-III). **b** Secondary structure and **c** 3D structural model of *E. coli* EF-P with selected boundaries between all domains (I-III) for hybrid construction (adapted from UniProt P0A6N4, and AlphaFold AF-P0A6N4-F1). Structure prediction and the confidence score (pLDDT) were calculated by AlphaFold. 3D model confidence: very high – in dark blue (pLDDT > 90); confident – in light blue (90 >pLDDT> 70); low – yellow (70 > pLDDT >50); very low – red (pLDDT < 50).

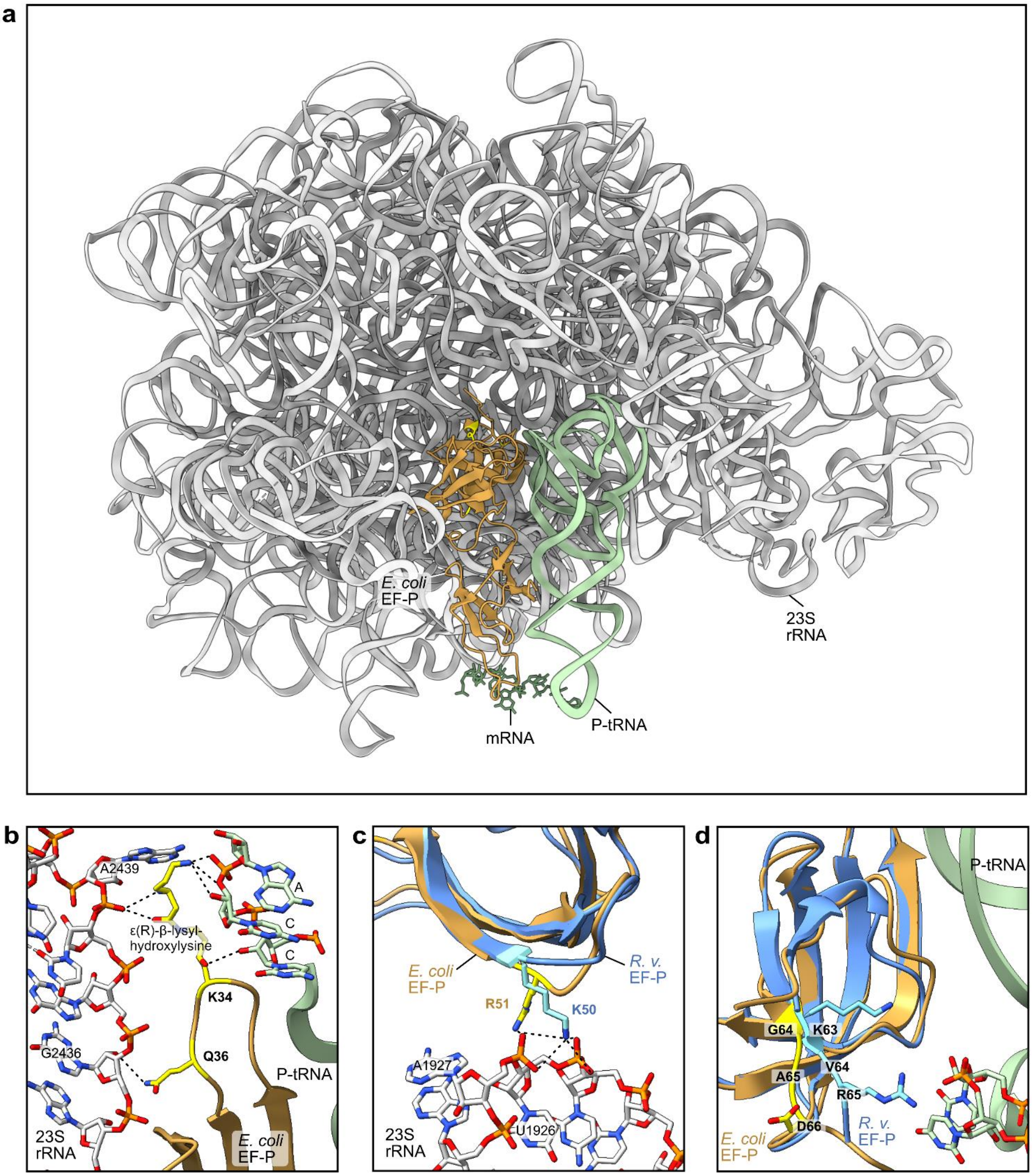
a
E. coli
EF-P
23S
rRNA
mRNA
P-tRNA
b
A2439
A
C
C
ε(R)-β-lysyl-
hydroxylysine
K34
Q36
G2436
P-tRNA
23S
rRNA
E. coli
EF-P
c
E. coli
EF-P
R. v.
EF-P
R51
K50
A1927
U1926
23S
rRNA
d
P-tRNA
G64
K63
A65
V64
R65
D66
E. coli
EF-P
R. v.
EF-P

**Supplementary Figure S6. Model of potential interactions between *E. coli* / *R. vannielii* EF-P and 23S rRNA / P-site-tRNA in context of amino acid substitutions P35Q, V50R, and 63-65_KVR in *S. venezuelae* EF-P**. The Cryo-EM structure of polyproline-stalled ribosome in the presence of *E. coli* EF-P was adopted from Huter *et al.*, 2017 [2] (PDB accession number: 6ENU). Structural models were generated using UCSF ChimeraX [3,4]. The *R. v.* EF-P structure was predicted using AlphaFold2 ColabFold (v1.5.2)[5,6]. **a** Schematic presentation of *E. coli* EF-P (brown), localized relative to the ribosome (23S rRNA) (grey), mRNA (dark green) and P-site-bound $\text{tRNA}^{\text{Pro}}$ (light green). **b** Potential interaction between Q36 of *E. coli* EF-P and 23S rRNA (G2436) is indicated with a dashed line. As described before [2] the interactions of ε(R)-ß-lysyl-hydroxylysine (after post translational modification of K34) with the CCA end of the P-site tRNA (P-tRNA) and the 23S rRNA (A2439) are shown in dashed lines. **c** Potential interactions between R51 of *E. coli* EF-P or K50 of *R. v.* EF-P with the phosphate backbone of the 23S rRNA (U1926, A1927), indicated in dashed lines. Domain I of the predicted *R. v.* EF-P was superimposed with domain I of the cryo-EM structure of *E. coli* EF-P. **d** Potential interactions between the motif KVR (positions 63-65 in *R. v.* EF-P) and the P-tRNA. For comparison, the position of the corresponding GAD motif (amino acids 64-66) in *E. coli* EF-P is marked. Amino acids K34, Q36, R51, G64, A65 and D66 in *E. coli* EF-P correspond to K33, P35, V50, T63, A64 and T65, respectively, in *S. venezuelae* EF-P (**Fig. 4a**).

### References


1. Conners, E.M., Davenport, E.J., and Bose, A. (2021). Revised Draft Genome Sequences of *Rhodomicrobium vannielii* ATCC 17100 and *Rhodomicrobium udaipurense* JA643. Microbiol Resour Announc *10*. 10.1128/MRA.00022-21.

2. Huter, P., Arenz, S., Bock, L.V., Graf, M., Frister, J.O., Heuer, A., Peil, L., Starosta, A.L., Wohlgemuth, I., Peske, F., et al. (2017). Structural Basis for Polyproline-Mediated Ribosome Stalling and Rescue by the Translation Elongation Factor EF-P. Molecular Cell *68*, 515-+. 10.1016/j.molcel.2017.10.014.

3. Goddard, T.D., Huang, C.C., Meng, E.C., Pettersen, E.F., Couch, G.S., Morris, J.H., and Ferrin, T.E. (2018). UCSF ChimeraX: Meeting modern challenges in visualization and analysis. Protein Sci *27*, 14-25. 10.1002/pro.3235.

4. Pettersen, E.F., Goddard, T.D., Huang, C.C., Meng, E.C., Couch, G.S., Croll, T.I., Morris, J.H., and Ferrin, T.E. (2021). UCSF ChimeraX: Structure visualization for researchers, educators, and developers. Protein Sci *30*, 70-82. 10.1002/pro.3943.

5. Jumper, J., Evans, R., Pritzel, A., Green, T., Figurnov, M., Ronneberger, O., Tunyasuvunakool, K., Bates, R., Zidek, A., Potapenko, A., et al. (2021). Highly accurate protein structure prediction with AlphaFold. Nature *596*, 583-+. 10.1038/s41586-021-03819-2.

6. Mirdita, M., Schutze, K., Moriwaki, Y., Heo, L., Ovchinnikov, S., and Steinegger, M. (2022). ColabFold: making protein folding accessible to all. Nat Methods *19*, 679-682. 10.1038/s41592-022-01488-1.
